# Organelle-selective click labeling coupled with flow cytometry allows high-throughput CRISPR screening of genes involved in phosphatidylcholine metabolism

**DOI:** 10.1101/2022.04.18.488621

**Authors:** Masaki Tsuchiya, Nobuhiko Tachibana, Kohjiro Nagao, Tomonori Tamura, Itaru Hamachi

## Abstract

Lipids comprise biomembranes and are involved in many crucial cell functions. While cellular lipid synthesis and transport appear to be governed by intricate protein networks, the whole scheme is insufficiently understood. Although functional genome-wide screening should contribute to deciphering the regulatory networks of lipid metabolism, technical challenges remain – especially for high-throughput readouts of lipid phenotypes. Here, we coupled organelle-selective click labeling of phosphatidylcholine (PC) with flow cytometry-based CRISPR screening technologies to convert organellar PC phenotypes into a simple fluorescence readout for genome-wide screening. This technique, named O-ClickFC, was successfully applied in genome-scale CRISPR-knockout screens to identify previously reported genes associated with PC synthesis (*PCYT1A, ACACA*), vesicular membrane trafficking (*SEC23B, RAB5C*), and non-vesicular transport (*PITPNB, STARD7*). Moreover, this work revealed previously uncharacterized roles of *FLVCR1* as a new choline transporter; *CHEK1* as a post-translational regulator of the PC-synthetic pathway, and *TMEM30A* as responsible for translocation of PC to the outside of the plasma membrane bilayer. These findings demonstrate the versatility of O-ClickFC as an unprecedented platform for genetic dissection of cellular lipid metabolism.

## Introduction

Lipids are a broad class of metabolites with diverse structures and myriad essential functions for the cell (Harayama and Riezman, 2018; Van Meer et al., 2008). Cellular lipid metabolism involves complicated protein networks that spatiotemporally regulate lipid synthesis and trafficking over various organelles (Vance, 2015). Thus, identifying enzymes/proteins in the network is a prerequisite for unravelling the molecular mechanisms underlying lipid-mediated biological processes, such as cell signaling and inter- or intra-organelle membrane transport.

Phosphatidylcholine (PC) is the most abundant phospholipid in eukaryotic membranes, and its metabolic abnormalities are associated with various diseases including cancer, lipodystrophy, muscular dystrophy, and retinal dystrophy (Glunde et al., 2011; Hoover-Fong et al., 2014; Mitsuhashi et al., 2011; Payne et al., 2014; Ridgway, 2013; van der Veen et al., 2017). Since the pioneering work by Kennedy and coworkers in 1956, significant progress has been made in understanding the enzymology of individual proteins involved in the PC biosynthetic pathway (Gibellini and Smith, 2010). However, whole protein networks regulating PC abundance and subcellular distribution remain elusive. Over the past four decades, functional genetic screens based on auxotrophy of yeast mutants have identified an extensive set of PC regulators (Atkinson et al., 1980). However, this conventional approach, which mostly relies on cell proliferation, has difficulty discovering genes that are not fatal but cause perturbations in PC dynamics. Fluorescent lipid-binding probes were used to identify the phospholipid scramblase from a cDNA library (Suzuki et al., 2013), but there is presently no fluorescent PC-binding probe. Although mass spectrometry (MS) coupled with subcellular fractionation can quantitatively analyze subcellular PC contents in a label-free manner (Schneiter et al., 1999), it is unsuitable for large-scale genomic screening due to its limited throughput. Until now, these technical challenges have limited our understanding of the link between PC metabolism and the 20,000 protein-coding genes in the human genome.

The recent emergence of genome-wide clustered regularly interspaced short palindromic repeats (CRISPR) knockout (KO) screens has the potential to provide a high-throughput platform for identifying key genetic factors in mammalian cells (Shalem et al., 2015). Pooled CRISPR screening builds on a library of cells bearing individual genetic perturbations. In a typical workflow, cells exhibiting the phenotype of interest are physically separated by cell viability or fluorescence-activated cell sorting (FACS), followed by next-generation sequencing (NGS) of sgRNA abundance to identify target genes. The pooled format makes it practical to handle large-scale libraries (∼20,000 genes for human) in a single experiment, thereby accelerating the discovery process of phenotype-to-genotype relationships. When this technology is applied to identify PC regulatory networks, the most critical task is deciding how to convert PC phenotypes into a simple readout for screening. Furthermore, with the aim of exploring key genes affecting subcellular/suborganelle PC distribution, the reporter requires organelle/suborganelle-level spatial resolution.

Recently, we reported a technology for selective labeling and imaging of PC in target organelles (Tamura et al., 2020). This method involves metabolic incorporation of azido-choline (N_3_-Cho) in live cells followed by an organelle-directed copper-free click reaction to modify the *de novo*-synthesized azido-PC (N_3_-PC) with fluorescent reporters. Using this method, PC in various organelle membranes can be simultaneously tagged with different-colored fluorescent dyes, which allows PC trafficking and dynamics between organelles to be observed in live cells by confocal microscopy.

Here, we developed a strategy to convert organelle lipid phenotypes into a simple fluorescence readout for genome-wide screening, named O-ClickFC (Organelle-selective Click chemistry coupled with Flow Cytometry). We selected several clickable compounds with diverse spectral properties that can specifically label PCs in the endoplasmic reticulum (ER)/Golgi apparatus, mitochondria, and outer leaflet of the plasma membrane (OPM), and established a workflow for FACS analysis with multiplexed compounds. Proof-of-principle experiments clearly demonstrated that this approach can distinguish defects in PC biosynthesis and organelle transport with flow cytometry. O-ClickFC was then successfully applied to CRISPR-KO screens, which identified a total of 52 genes involved in PC metabolism and trafficking, including known genes *PCYT1A, ACACA*, and *STARD7*. More significantly, we discovered previously unrecognized roles for *CHEK1, FLVCR1*, and *TMEM30A* in PC biosynthesis, choline transport, and cell-surface PC translocation, respectively.

## Results

### Development of O-ClickFC

O-ClickFC comprises three steps (Figure 1): (1) metabolic incorporation of an N_3_-Cho tracer into intracellular PC via endogenous pathways; (2) spatially selective labeling of N_3_-PC with organelle-targeting clickable dyes (OCDs); and (3) flow-cytometric measurements of fluorescently labeled lipids in target organelles. Subcellular N_3_-PC distribution can be spatiotemporally fixed by rapid and multiplexed organelle-selective labeling with different fluorescence wavelengths. The abundances of N_3_-PC in target organelles can be converted into fluorescence intensities of the corresponding OCDs, which are measurable with FACS. Thus, our method can readily provide informative snapshots of local N_3_-PC quantity, which allows interpretation of whether the phenotype occurs due to global abnormalities in PC synthesis or subcellular transport disorders (Figure 1).

**Figure 1.**
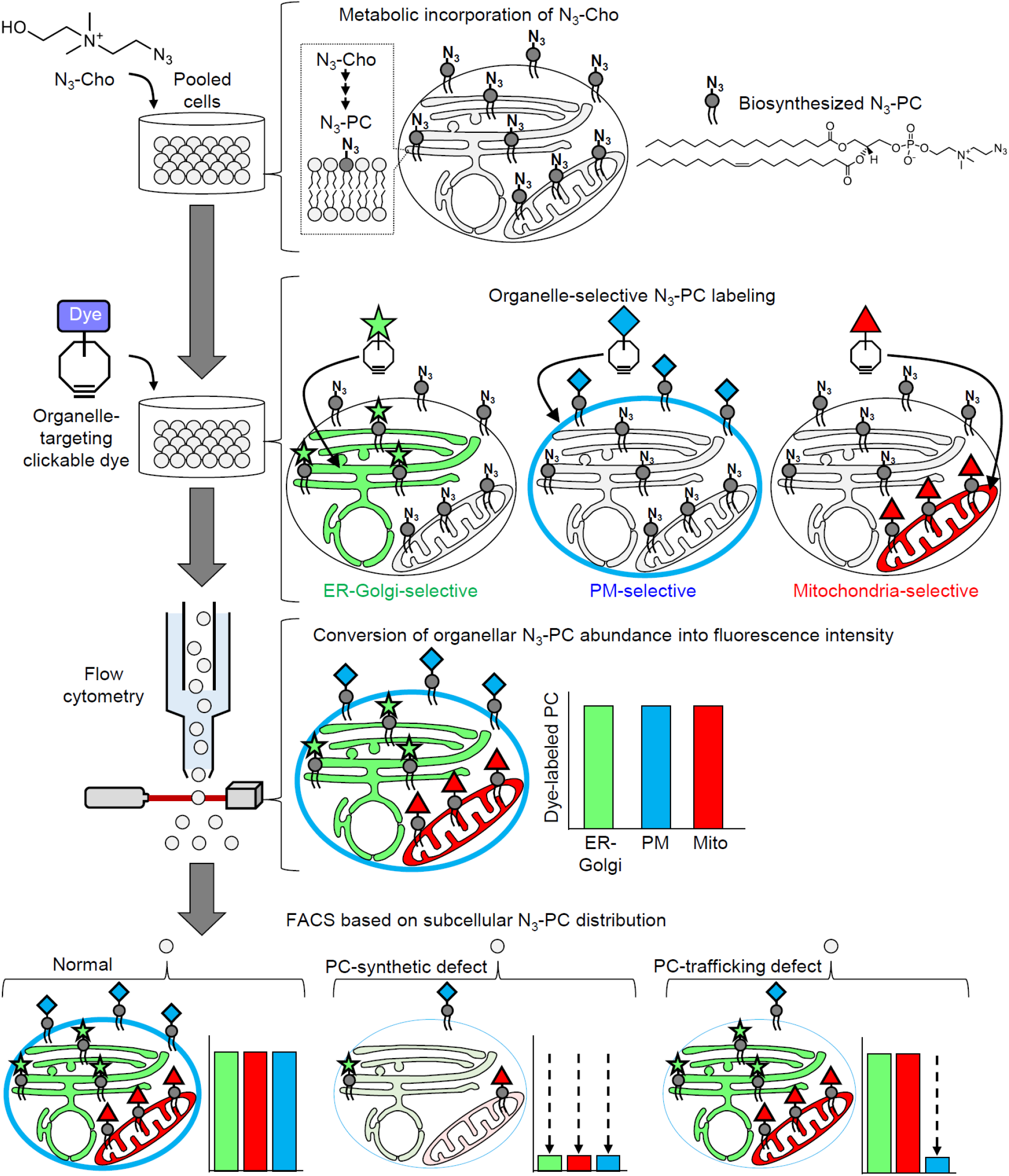
Workflow for flow cytometric analysis of subcellular PC distribution. N_3_-Cho is metabolically incorporated in cells to produce N_3_-PC via the PC-synthetic pathway. The *de novo* synthesized N_3_-PC is distributed to various organelles. Organelle-targeting clickable dyes (OCDs) selectively react with N_3_-PC to generate dye-labeled PC in the corresponding subcellular compartments, allowing information about the localization and quantity of N_3_-PC to be obtained from fluorescence wavelength and intensity, respectively. Thus, on the basis of changes in fluorescence patterns of dye-labeled PC, single cells exhibiting specific subcellular lipid phenotypes can be identified and isolated with FACS.

O-ClickFC requires a labeling signal with high dynamic range, as well as a suitable combination of OCDs without overlapping wavelengths in FACS-based quantitative analysis. To identify the optimal OCDs, we employed nine clickable dyes with diverse spectral properties (Figure S1A) and tested their labeling capability for organelle PCs with confocal microscopy and FACS. K562 human leukemic cells, which have been widely used in genetic screening studies (To et al., 2019; Wang et al., 2015), were incubated with 10 μM N_3_-Cho for 1 day followed by treatment with OCDs for 15–30 min. Imaging analysis with confocal microscopy showed that, in addition to our original Rhodol-DBCO, the hydrophobic BODIPY derivatives BDP and 8AB selectively localized to the ER/Golgi and labeled N_3_-PC present there (Figure 2A and S1B,C). In line with our previous report (Tamura et al., 2020), cationic dyes such as RhodB, Cy3, and Cy5 derivatives selectively labeled mitochondrial PC, while membrane-impermeable Alexa Fluor (AF) 405, AF488, and AF647 derivatives exclusively visualized PC in the OPM (Figure 2A and S1B,C). Flow cytometry revealed that fluorescence intensities of N_3_-Cho-treated cells were higher than those of non-treated cells for all OCDs (Figure 2B and S2A). The fold-change ratio, depending on the labeling efficiency and background signal from unreacted reagents after washing cells, varied with each dye and the experimental conditions (Figure 2B and S2A). We decided to use the following OCDs for single labeling because they showed greater fold-changes under optimal conditions (Figure S2A–C): BDP or 8AB for the ER-Golgi, Cy3 for mitochondria, and AF405 or AF647 for the OPM. For multiplexed labeling, we chose a combination of OCDs with no spectral overlap and the highest signal-to-noise ratios [e.g. blue-emitting 8AB (Ex. 394 nm/Em. 458 nm) and red-emitting Cy3 (Ex. 555 nm/Em. 564 nm) for ER-Golgi and mitochondrial labeling, respectively] (Figure S2D).

**Figure 2.**
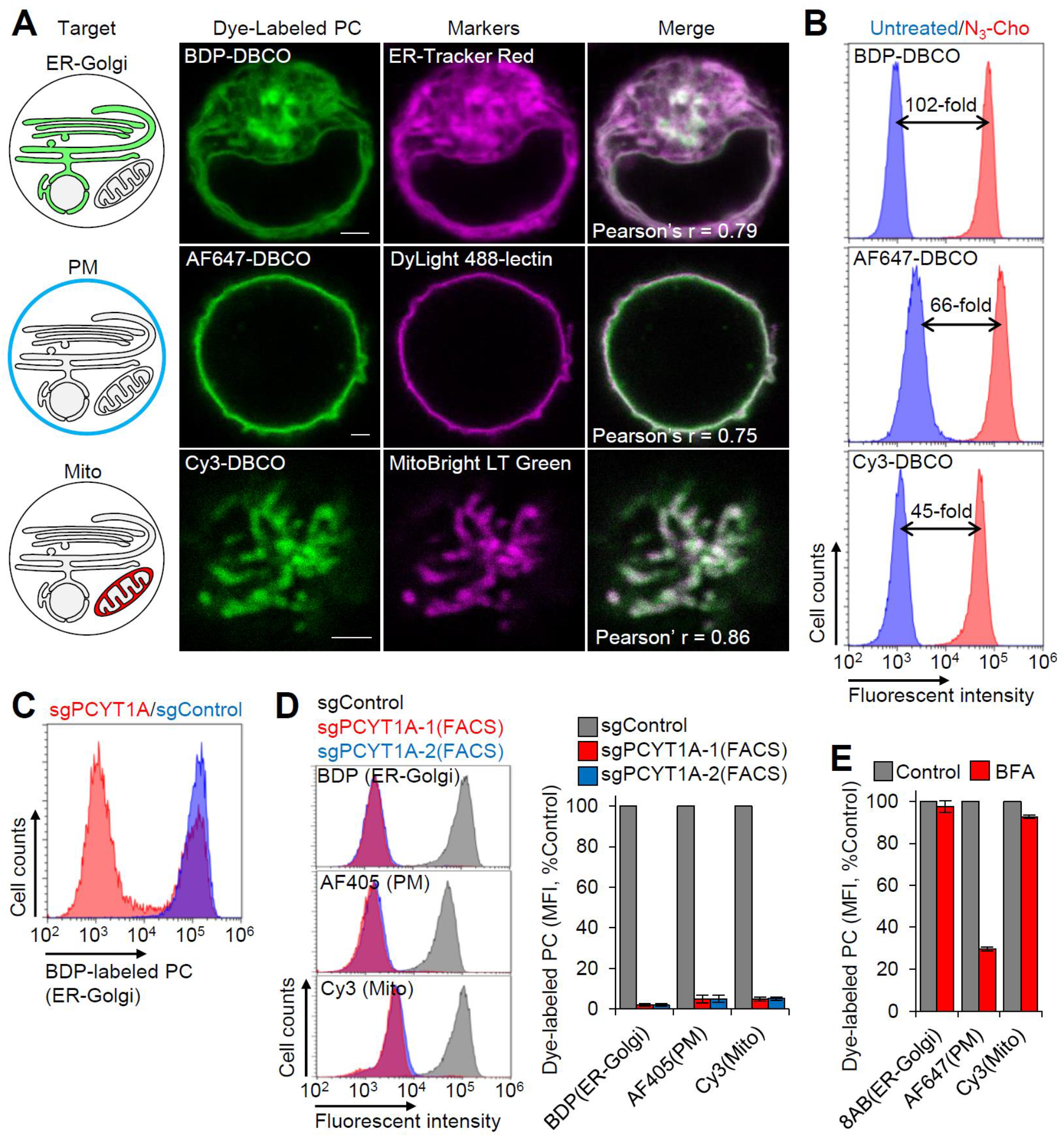
Flow cytometric measurements of N_3_-PC abundance at the organelle level. (A) Confocal imaging of organelle-selective N_3_-PC labeling. N_3_-Cho-treated K562 cells were labeled with ER-Golgi-targeting BDP-DBCO, PM-targeting AF647-DBCO, and mitochondria-targeting Cy3-DBCO. Corresponding organelle markers were used for colocalization analysis by confocal microscopy. Scale bars represent 2 μm. (B) Quantitative measurements of organelle-selective N_3_-PC labeling. Non-treated or N_3_-Cho-treated K562 cells were labeled with BDP-DBCO, AF647-DBCO, and Cy3-DBCO, and analyzed by flow cytometry. Median fluorescence intensity (MFI) was used to calculate fold changes between the two groups. (C) K562 cells transduced with sgControl and sgPCYT1A were labeled with BDP-DBCO, followed by analysis with flow cytometry. The majority of sgPCYT1A-transduced cells displayed a reduction in BDP fluorescence. (D) Labeling pattern of cells with PC synthesis defects. FACS-isolated *PCYT1A*-KO cells [the dimmer subpopulation in (C)] were treated with N_3_-Cho and labeled with BDP (ER-Golgi), AF405 (PM), and Cy3 (mitochondria). MFIs analyzed by flow cytometry were used to compare N_3_-PC labeling between control and *PCYT1A*-KO cells (n = 3). (E) Labeling pattern of cells with BFA-mediated PC transport defects. K562 cells were treated with N_3_-Cho in the presence of 10 μM BFA for 5 hours and labeled with 8AB (ER-Golgi), AF647 (PM), and Cy3 (mitochondria), followed by flow cytometry (n = 3). Bar graphs represent mean ± SEM.

To evaluate the quantitative cell-separation performance of multiplexed labeling, we prepared a mixed population of cells preincubated with varying N_3_-Cho concentrations (0, 0.1, 1, 10 μM) and carried out dual N_3_-PC labeling for the ER-Golgi and another organelle with corresponding OCDs. Two-color flow cytometric analysis correctly divided the bulk population into four groups with different fluorescence levels corresponding to each N_3_-Cho concentration (Figure S2E). These data demonstrate that O-ClickFC can separate cells on the basis of intracellular N_3_-PC distribution and abundance with a sufficient dynamic range (∼1,000 in flow cytometric quantification).

We next examined whether this method could reveal subcellular phenotypes associated with genetic perturbations of PC biosynthesis. The major PC biosynthetic pathway in mammalian cells, the CDP-choline pathway, comprises three steps: (1) phosphorylation of choline catalyzed by choline kinase (CK); (2) conversion of phosphocholine to CDP-choline by CTP: phosphocholine cytidylyltransferase (CCT); and (3) transfer of CDP-choline to diacylglycerol by choline phosphotransferase (CPT) in the ER and Golgi apparatus (Gibellini and Smith, 2010) (Figure S3A). CCT is the rate-limiting enzyme of this pathway, and suppression of CCTα (a predominant isoform of CCT) can decrease intracellular PC to inhibit cell growth (Cornell and Ridgway, 2015; Esko and Raetz, 1980). Thus, we transduced a bulk population of Cas9-expressing K562 cells with sgRNAs of *PCYT1A* encoding CCTα (Figure S3A) and evaluated alterations in amounts of *de novo*-synthesized N_3_-PC using O-ClickFC. After ER-Golgi-selective labeling, sgPCYT1A-transduced cells were divided into two populations (exhibiting bright and dim fluorescence) by flow cytometry, while control cells (without sgPCYT1A-transduction) exhibited a single bright population (Figure 2C). The population exhibiting weak fluorescence was collected by FACS and expressed a characteristic phenotype of *PCYT1A*-KO, namely, suppressed cell proliferation and synthetic defects in both natural form-and N_3_-PC (DeLong et al., 1999) (Figure S3B-E). The same experiments were performed with mitochondria-and OPM-selective OCDs, which also showed reduced fluorescence intensity in *PCYT1A*-KO cells (Figure 2D). These results, which are consistent with previous reports of *PCYT1A*-KO causing a global decrease in PC contents of all subcellular membranes (Andrejeva et al., 2020; Esko and Raetz, 1980), successfully demonstrate the feasibility of O-ClickFC for detection of genotype-to-phenotype correlation of PC synthesis defects.

We subsequently examined whether this FACS-based analysis could readout PC trafficking defects. K562 cells were incubated for 5 hours with N_3_-Cho in the presence of brefeldin A (BFA), an inhibitor of Golgi-mediated anterograde membrane trafficking from the ER to plasma membrane (Wood et al., 1991), followed by labeling with each OCD. Fluorescence intensities of cells labeled by ER-Golgi and mitochondria OCDs were not significantly changed with BFA treatment (within ± 10% of control) (Figure 2E). In stark contrast, when labeled with the OPM-staining dye, the fluorescence signal dropped by ∼70% in both flow cytometry and microscopic observations of individual cells (Figure 2E and S3F), in agreement with the phenotype of BFA-induced impairment of PC transport. Notably, duplexed labeling with ER-Golgi- and OPM-OCDs yielded the same result (Figure S3G).

Overall, these data prove that O-ClickFC can discern and enrich cells showing deficiencies in PC synthesis and trafficking on the basis of differences in the fluorescence intensity of each organelle.

### Genome-scale pooled CRISPR-KO screens focusing on PC biosynthesis

Having confirmed the potential ability of O-ClickFC, we sought to apply it for genome-wide CRISPR-KO screening to identify genes associated with PC biosynthesis. In our initial trial, Cas9-expressing K562 cells were transduced with the GeCKOv2 library (19,050 genes, 65,383 sgRNAs, 1–3 guides/gene) (Joung et al., 2017; Sanjana et al., 2014) followed by labeling with ER-Golgi OCD (BDP-DBCO) (Figure 3A). The dimmest 1% of cells were collected by FACS and subjected to NGS analysis to rank genes according to sgRNA abundance ratio (sort/unsort) (Figure 3A,B and S4A). *PCYT1A* ranked 61st (top 0.3%), indicating enrichment of genes related to PC synthesis in the collected cell population. To evaluate the validity of our screens, we selected 22 genes with known roles or potential contributions in PC synthesis (e.g., kinases, signaling proteins, and transcriptional regulators) from the top 100 genes and conducted PC labeling with cells transduced by a corresponding individual sgRNA. Here, we defined a hit gene as a gene whose knockout reduced PC labeling intensity by more than 20% in the individual validation, and four genes (*PCYT1A, CHEK1, GOLGA8O*, and *AGAP4*) were verified to meet this criterion (Figure S4B). The true positive rate was low (18%) using the GeCKO v2 library, but was improved when using the more recently established Brunello library (19,114 genes, 76,441 sgRNAs): 33 out of 77 genes were assigned as hits (43% of the true positive rate) (Figure S4C). We thus decided to use the Brunello library for subsequent CRISPR-KO screening.

**Figure 3.**
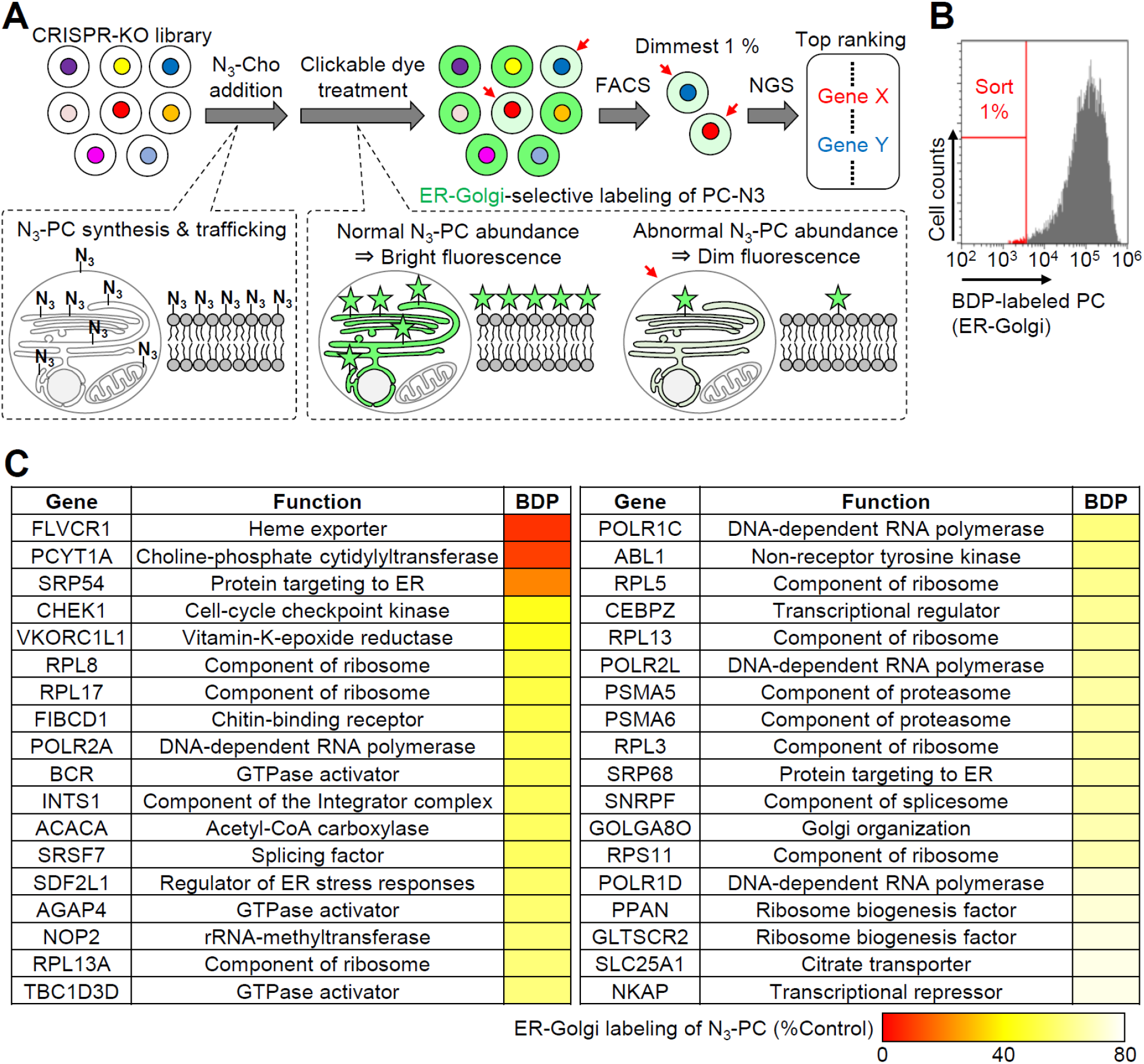
O-ClickFC-based CRISPR-KO screening to identify genes involved in PC biosynthesis. (A) Experimental workflow. Cas9-expressing K562 cells were transduced with a genome-wide lentiviral sgRNA library, cultured with N_3_-Cho for 1 day, and labeled with ER-Golgi-selective OCD. Target cells with reduced abundance of labeled PC (red arrows) were sorted by FACS and subjected to NGS analysis to produce a ranked list of candidate genes. (B) GeCKOv2 library-expressing K562 cells with decreased fluorescence (1% of the population) were sorted (FACS gating strategy in Figure S4A). (C) A total of 36 hit genes identified from GeCKOv2 and Brunello screens. The criteria of hits were set as follows: two or more distinct sgRNAs targeting the same gene displayed < 80% ER-Golgi N_3_-PC compared with a nontargeting control sgRNA. The average from multiple sgRNAs is represented by a pseudocolor scale.

The combined 36 hits screened from GeCKO v2 and Brunello libraries are shown in Figure 3C. In addition to *PCYT1A*, two known contributors to PC synthesis, *ACACA* (acetyl CoA carboxylase 1) and *SLC25A1* (citrate transport protein), were found (Figure 3C). These results indicate that O-ClickFC can also identify genes involved in fatty acid synthesis. To date, the relevance of the other 33 genes for PC synthesis is unknown. Given that CCTα activity is reversibly regulated by phosphorylation/dephosphorylation at its C-terminal serine-proline sites during the cell cycle, some sort of signaling pathways should be involved in regulation of CCTα activity; at present, these pathways remain unclear. Of the hit genes discovered in our screening, we were interested in *CHEK1* (coding checkpoint kinase 1, CHK1) (Figure 3C and S4B), which controls cell cycle via the CDC25A–cyclin-dependent kinase 2 (CDK2) pathway (Zhang and Hunter, 2014), and we hypothesized that it might control CCTα activity. In our experiments, we indeed found that PC labeling intensity was decreased by a CHK1 inhibitor (CHK1i) but was rescued by a CDK2 inhibitor (CDK2i), and hypothesized that it might control CCTα activity. In our experiments, we indeed found that, the PC labelling intensity was decreased by CHK1 inhibitors (CHK1i) but it was rescued by CDK2 inhibitor (CDK2i) (Jorda et al., 2018; Schuler et al., 2017) (Figure 4A). The similar effect was also observed in the *de novo* synthesis of native-form PC (Figure S5A), indicating the CHK1-CDC25A-CDK2 pathway is associated with PC synthesis. We next examined whether this pathway regulates the phosphorylation states of CCTα. Phos-tag electrophoresis (Kinoshita et al., 2006) (Weinhold et al., 1994; Yue et al., 2020). Upon CHK1i treatment, the CCTα phosphorylation level in choline-free medium was significantly increased (Figure 4B, lane 2 and 3), while coincubation of CDK2i with CHK1i attenuated this effect (Figure 4B, lane 3 and 4). These data clearly reveal the regulation of CCTα phosphorylation states by the CHK1-CDC25A-CDK2 pathway (Figure 4C).

**Figure 4.**
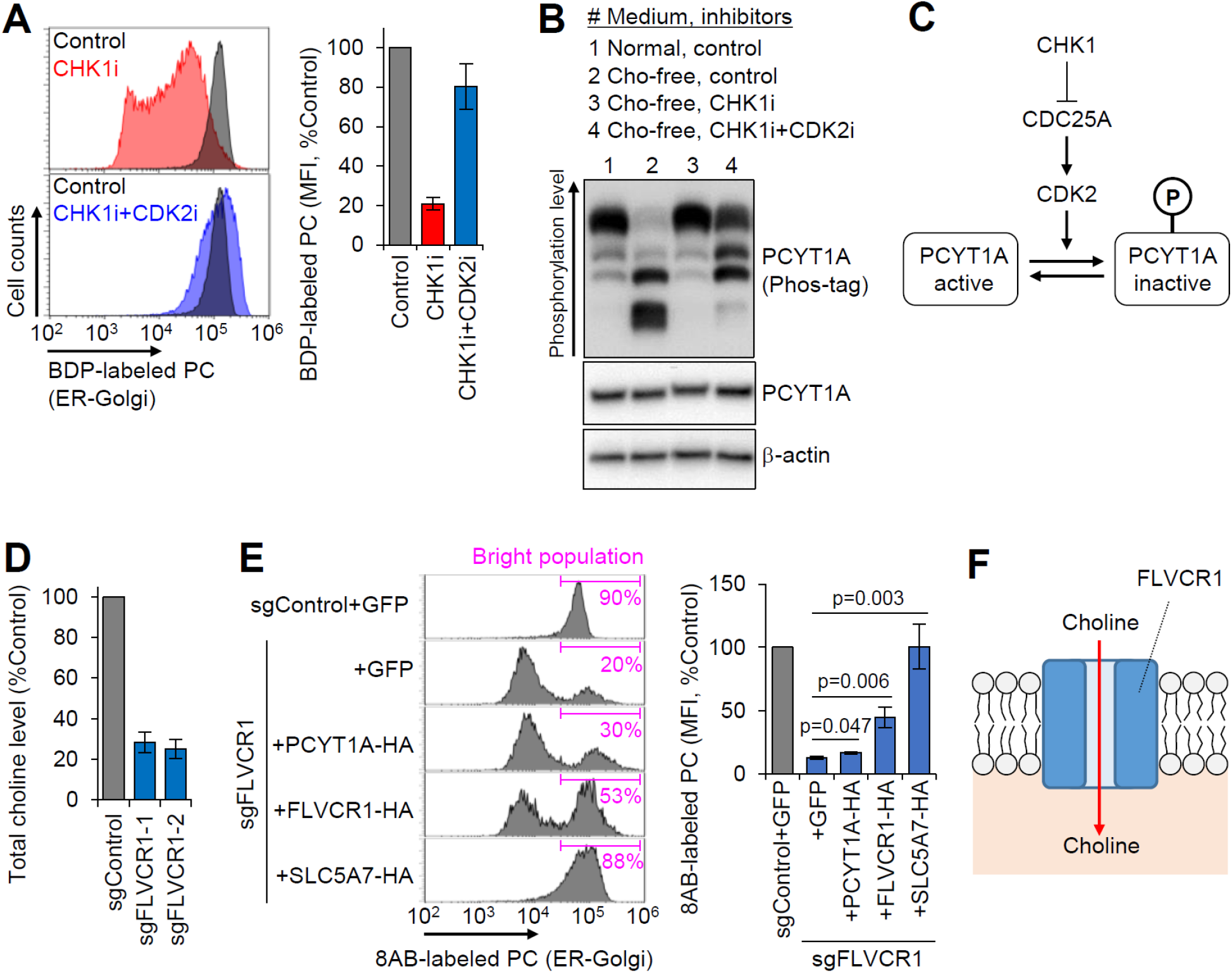
Roles of *CHEK1* and *FLVCR1* in PC synthesis. (A-C) CHK1 regulates PC synthesis via modulating phosphorylation state of PCYT1A. (A) Flow cytometric analysis of BDP-labeled PC in K562 cells treated with CHK1 inhibitor (PF-477736) and CDK2 inhibitor (Cdk2 inhibitor II) (n = 3–10). The inhibitory effect of CHK1i on N_3_-PC synthesis was counteracted by CDK2i. (B) Effect of CHK1 and CDK2 inhibition on PCYT1A phosphorylation was analyzed by phos-tag electrophoresis, wherein highly phosphorylated PCYT1A migrated slower than its poorly phosphorylated form. Choline depletion promoted dephosphorylation of PCYT1A (lane 1 vs 2). Inhibition of CHK1 in choline-free culture provoked PCYT1A to hyper-phosphorylate state (lane 2 vs 3). Dual treatment of CHK1 and CDK2 inhibitors led to mild phosphorylation of PCYT1A (lane 3 vs 4). (C) Schematic showing how the CHK1-CDC25A-CDK2 pathway regulates PCYT1A-mediated PC synthesis. (D–F) FLVCR1 acts as a choline transporter. (D) Endogenous choline levels in control-and *FLVCR1*-KO K562 cells (n = 3). (E) The rescue effect of exogenous SLC5A7 expression on ER-Golgi N_3_-PC level in *FLVCR1*-KO cells. *FLVCR1*-KO K562 cells were infected with lentiviruses expressing GFP (control), HA-tagged *PCYT1A, FLVCR1*, or *SLC5A7*, and subjected to flow cytometric analysis of ER-Golgi N_3_-PC (n = 3–6, Student’s t-test). (F) Schematic illustration showing FLVCR1-mediated choline transport. Bar graphs represent mean ± SEM.

*FLVCR1* was also selected for the follow-up study because its deficiency caused a strong reduction in PC labeling and cell growth inhibition, comparable with those of *PCYT1A*-KO (Figure 3C and S5C). *FLVCR1*, a member of the major facilitator superfamily (MFS), was previously identified as a plasma membrane heme exporter (Quigley et al., 2004), but its function related to PC metabolism has never been reported. MFS transporters are generally considered to transport a wide range of substrates across membranes by uniport, symport, or antiport (Quistgaard et al., 2016). Therefore, we assumed that *FLVCR1* may play an additional role as a choline importer. To confirm this, we first quantified natural choline extracted from cytoplasm using a choline oxidase-mediated coupled enzyme assay, which revealed that the loss of *FLVCR1* dramatically reduced the amount of endogenous choline inside cells (by nearly 80%) (Figure 4D). Note that these phenotypes of *FLVCR*-KO (decrease in PC synthesis and choline uptake) are the same phenotypes associated with major choline transporter inhibition (Taylor et al., 2021). Moreover, we found that the lower PC labeling intensity in *FLVCR1*-KO cells was partially restored by re-expression of *FLVCR1* and completely rescued by overexpression of *SLC5A7* (a high affinity choline transporter specifically expressed in cholinergic neurons) (Okuda et al., 2000); whereas overexpression of *PCYT1A* had little effect (Figure 4E). Moreover, recoveries in choline uptake and cell proliferation were observed in *SLC5A7*-expressing *FLVCR1*-KO cells (Figure S5D,E). These data clearly demonstrate that *FLVCR1* acts as a choline transporter in K562 cells (Figure 4F).

### Genome-scale pooled CRISPR screens focusing on subcellular PC trafficking

We finally applied O-ClickFC to identify genes associated with subcellular PC distribution. For screening focused on PC transport from the ER-Golgi to the OPM, we performed duplex labeling with two OCDs (BDP-DBCO and AF405-DBCO, respectively) in K562 cells transduced by the Brunello library (Figure 5A). Two-color flow cytometry showed the presence of a small cell population with decreased labeling intensity in the OPM, but that maintained labeling in the ER-Golgi (Figure 5B and S6A), suggesting impaired PC trafficking to the OPM. This population was clearly distinguishable from the majority population exhibiting an unchanged or decreased PC content in both the ER-Golgi and OPM. We collected the minor cell population and subjected it to the same duplex labeling-based FACS sorting again to increase selection pressure. The NGS of collected cells and gene ontology analysis indicated enrichment of genes related to intracellular transport in the top 100 ranked genes. From the top 100, we selected 28 genes known or unknown to be involved in membrane transport and conducted duplex labeling in cells with individual knockouts of the selected genes. Twelve genes satisfied the following criteria: < 80% OPM-labeling and > 60% ER-Golgi labeling, most of which (eight genes) have functions related to vesicle transport and lipid transfer (Figure 5C and S6B).

**Figure 5.**
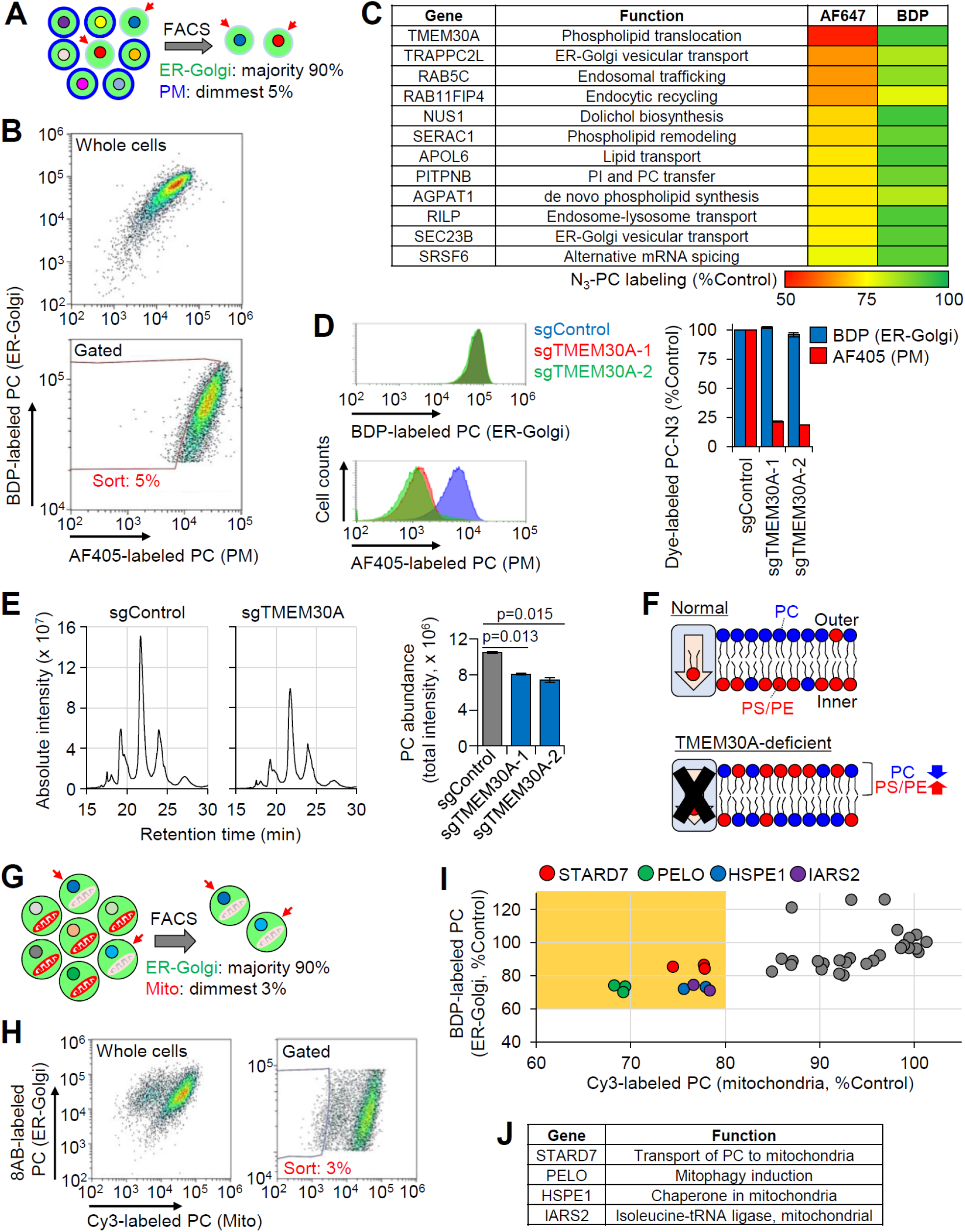
O-ClickFC-based CRISPR-KO screening focusing on N_3_-PCsubcellular PC trafficking. (A) CRISPR-KO screen based on N_3_-PC distribution in the ER-Golgi and PM. K562 cells transduced with a genome-wide CRISPR-KO library were labeled with ER-Golgi-selective OCD (BDP) and PM-selective OCD (AF405). Target cells with normal BDP fluorescence and weak AF405 fluorescence were sorted by FACS. (B) Brunello library-expressing K562 cells with decreased PM N_3_-PC (∼5% of the parent population) were sorted (FACS gating strategy in Figure S6A). (C) List of 12 hit genes presumed to be involved in PC transport from the ER-Golgi to the PM. Heatmap shows averaged N_3_-PC levels in PM (AF647) and ER-Golgi (BDP) for each gene (data from Figure S6B). (D-F) TMEM30A allows PC translocation to the outer leaflet of PM (OPM). (D) N_3_-PC levels in ER-Golgi and PM in control and *TMEM30A*-KO cells (n = 3). (E) LC-MS/MS analysis of endogenous PC extracted from OPM of control and *TMEM30A*-KO cells by methyl-α-cyclodextrin-mediated lipid exchange (n = 2, Student’s t-test). (F) Schematic illustration showing decreased PC and increased PS/PE in the OPM of TMEM30A-deficient cells. (G) CRISPR-KO screen based on N_3_-PC distribution in the ER-Golgi and mitochondria. Target cells with normal BDP fluorescence and weak Cy3 fluorescence were sorted by FACS. (H) Brunello library-expressing K562 cells with decreased mitochondrial N_3_-PC (∼3% of the parent population) were sorted (FACS gating strategy in Figure S6G). (I) Individual testing of 37 sgRNAs targeting 30 genes that were selected from the top 200. Two-dimensional dot plot displays fluorescence levels of Cy3-labeled mitochondrial PC and BDP-labeled ER-Golgi PC in sgRNA-transduced K562 cells (represented as mean). Multiple sgRNAs targeting *STARD7, PELO, HSPE1*, and *IARS2* showed > 60% ER-Golgi N3-PC and < 80% mitochondrial N_3_-PC compared with a nontargeting sgRNA (orange area). (J) List of four hit genes presumed to be involved in PC transport from the ER-Golgi to mitochondria. Bar graphs represent mean ± SEM.

Among them, we focused on *TMEM30A* (CDC50A), whose sgRNAs caused the lowest OPM labeling intensity (Figure 5D). In the mammalian plasma membrane, PC primarily resides in the outer leaflet, while the majority of aminophospholipids, such as phosphatidylserine (PS) and phosphatidylethanolamine (PE), are localized in the inner leaflet (Van Meer et al., 2008; Murate et al., 2015). *TMEM30A* is known to encode an essential subunit of aminophospholipid flippases that translocate PS and PE from the outer to the inner leaflet (Andersen et al., 2016; Kato et al., 2013; Segawa et al., 2014). However, it has not been reported whether the loss of *TMEM30A* function affects PC distribution in the plasma membrane bilayer, although the literature suggests an involvement of *TMEM30A* in PC flipping (Segawa et al., 2014). Thus, we investigated whether the amount of natural-form PC is reduced in the OPM of *TMEM30A*-KO cells. As with previous reports (Kato et al., 2013; Segawa et al., 2014), increased amounts of cell surface PS and PE in *TMEM30A*-KO K562 cells were observed by Annexin V and Duramycin staining (Fu et al., 2021; Suzuki et al., 2013) respectively, confirming that the phenotype is reproduced in our experimental conditions (Figure S6C). We subsequently performed quantitative liquid chromatography-mass spectrometry (LC-MS) analysis combined with established membrane fractionation methods to determine PC amounts in the plasma membrane. K562 cells were incubated with a deuterated choline (choline-D9), followed by extraction of lipids from whole-cell lysate or the isolated plasma membrane bilayer. In both of these fractions, we observed negligible differences in the amount of isotope-labeled PC (PC-D9) between *TMEM30A*-KO and control cells (Figure S6D,E). These results indicate that *TMEM30A*-KO impacts neither PC biosynthesis nor total PC contents in the plasma membrane bilayer. We next evaluated the quantity of endogenous PC in the OPM by selectively extracting phospholipids in the OPM by methyl-α-cyclodextrin, in accordance with previous reports (Li et al., 2016). LC-MS showed a substantial decrease in endogenous PC in the OPM fraction of *TMEM30A*-KO cells compared with control cells (Figure 5E and S6F). Overall, these data reveal that *TMEM30A*-KO downregulates cell surface PC amounts, suggesting a contribution of *TMEM30A* to proper asymmetric PC distribution in the plasma membrane bilayer (Figure 5F).

To screen for genes implicated in mitochondrial PC transport, OCDs for the ER-Golgi and mitochondria (8AB-DBCO and Cy3-DBCO, respectively) were concurrently employed (Figure 5G). Similar to the results described above, two-dimensional flow cytometry data successfully discriminated a subpopulation emitting lower fluorescence intensity in mitochondria while maintaining it in the ER-Golgi (Figure 5H and S6G). NGS and subsequent validation using individual sgRNAs (30 genes from the top 200 ranked genes) identified four hit genes (*STARD7, PELO, HSPE1*, and *IARS2*) as potential regulators of mitochondrial PC transport (Figure 5I,J). It should be noted that *STARD7*, encoding mitochondrial StAR-related lipid transfer protein 7, was previously reported to specifically transfer PC from the ER to mitochondria via a non-vesicular system (Horibata and Sugimoto, 2010; Horibata et al., 2016; Wong et al., 2019). This result is evidence that O-ClickFC can identify genes relevant to PC transport through both vesicular transport (a major PC trafficking pathway) and monomeric diffusion mediated by PC-specific transfer proteins.

## Discussion

We combined organelle-limited click labeling of phospholipids with flow cytometry to develop O-ClickFC to identify human genes involved in PC metabolism. O-ClickFC can directly measure the abundance of *de novo*-synthesized PC by FACS at organelle resolution, which allows high-throughput CRISPR screening focused on subcellular PC distribution. Compared with recent examples of FACS-based CRISPR screens using lipid metabolism reporters such as fluorescent lipid (cholesterol)-binding proteins (Lu et al., 2022), phospholipase D activity-based click labeling (Bumpus et al., 2021), or a fluorescent PC analog (Maruoka et al., 2021), O-ClickFC offers a unique and unprecedented platform to elucidate uncertain links between PC metabolism and the whole genome.

Several conclusions can be drawn from our O-ClickFC based CRISPR screens. Three genes known to contribute to synthesis of PC intermediate metabolites (*PCYT1A, ACACA*, and *SLC25A1*) were identified, demonstrating that the screen with a single ER-Golgi-OCD enabled capture of genes associated with the PC-synthetic pathway. We also found that the majority of hit genes identified as PC synthesis regulators are involved in fundamental cellular processes such as gene expression, cell cycle, and protein folding (e.g., *SRP54, RPL8*, and *POLR2A*; Figure 3C). Given that PC metabolism is dynamically modulated during the cell cycle (Jackowski, 1994), knockouts of these genes may impact on PC synthesis. Five genes of 12 hits obtained from the screen with ER-Golgi and OPM-OCDs are related to intracellular vesicular transport pathways (*TRAPPC2L, RAB5C, RAB11FIP4, RILP*, and *SEC23B*; Figure 5C). This result is consistent with the general understanding that newly synthesized PC is transported from the ER-Golgi to the plasma membrane via vesicular secretion (Van Meer et al., 2008), thus representing the robustness of O-ClickFC-based screens for PC-transport related genes. A potential limitation of O-ClickFC is that the steric hindrance of the methylazide group of N_3_-PC might prevent biological processes mediated by critical interactions between the choline head group of PC and proteins. However, we successfully captured *PITPNB* and *STARD7* (Figure 5C,J), both of which recognize the choline head group of PC and selectively transport it between organelle membranes (Ashlin et al., 2021; Haberkant et al., 2013; Horibata and Sugimoto, 2010; Vordtriede et al., 2005). Detection of these two hits indicates that the biological impact of azido group incorporation is minimal or negligible. O-ClickFC is thus feasible for screening of genes involved in non-vesicular lipid transport, as well as vesicular transport. Another concern is that the organelle selectivity of labeling by OCDs potentially depends on the physiological states of subcellular membranes to some extent, which may provide false positive/negative results – particularly for mutant cells. However, this issue can be readily verified by imaging experiments with confocal microscopy (Figure S3F), demonstrating the flexibility of O-ClickFC in terms of compatibility with both FACS and microscopic analyses. It should also be noted that genes specifically associated with sphingomyelin metabolism were not obtained in this work, even though the metabolic incorporation of azido-choline covers all choline-containing phospholipids (Jao et al., 2015). This could be explained by sphingomyelin being less abundant than PC throughout subcellular membranes (Van Meer et al., 2008), meaning fluctuations in its abundance have a smaller effect on labeling signal intensity, which is in line with our previous study (Tamura et al., 2020).

Among hit genes whose relationship to PC biosynthesis/transport was not known, we unveiled unexpected roles of *FLVCR1, CHEK1*, and *TMEM30A* in PC metabolism. *FLVCR1* has been identified as a heme exporter and has thereafter been studied in the context of heme metabolism (Khan and Quigley, 2013; Quigley et al., 2004). Here, we discovered an additional role of *FLVCR1* as a choline transporter. Choline uptake via choline transporters is essential for the survival of mammalian cells (Holmes-Mcnary et al., 1997). *SLC44A2*, a known choline transporter (Traiffort et al., 2013), is constitutively expressed in K562 cells (To et al., 2019), but is not required for survival (Wang et al., 2015). A set of our experimental data revealed that *FLVCR1* encodes a major transporter for the choline-uptake pathway of K562 cells and, therefore, is essential for cell proliferation (Figure 4D and S5C). *FLVCR1* is reportedly expressed in many cell types and is often associated with human disease (Quigley et al., 2004; Rajadhyaksha et al., 2010). Our findings indicate an interesting possibility that *FLVCR1* may play a broad role in PC metabolism. CHK1 regulates cell cycle progression by phosphorylating its downstream targets (Zhang and Hunter, 2014). We demonstrated that the CHK1-CDC25A-CDK2 pathway regulates PC synthesis via PCYT1A phosphorylation. It was previously assumed that the carboxyl-terminal domain of PCYT1A is a potential substrate of proline-directed kinases such as CDKs (Cornell and Ridgway, 2015), which agrees well with our present results. Given that pharmacological inhibition of PC synthesis is an intriguing approach for cancer treatment (Glunde et al., 2011), our finding may open a new avenue to suppress malignant cell growth by CHK1-mediated regulation of PCYT1A activity. TMEM30A is regarded as critical for establishing the asymmetric distribution of PS but not PC (Andersen et al., 2016). We found, by using O-ClickFC, that TMEM30A could play an additional role in translocation of PC to the outer leaflet of the plasma membrane. A recent report suggested that a neuronal disease-associated mutation in flippase *ATP11A*, whose activities are regulated by TMEM30A, causes aberrant PC flipping and decreases the concentration of PC in the OPM (Segawa et al., 2021). Combined with our result, it may be proposed that malfunctions of TMEM30A cause abnormal PC distribution in the plasma membrane bilayer, leading to disorders such as neurological deterioration.

We expect that the O-ClickFC approach can easily be expanded to evaluate a variety of lipids, including glycerophospholipids, sphingolipids, and inositol lipids, by combination with the established metabolic incorporation of an azido group into these lipids (Garrido et al., 2015; Greaves et al., 2017; Neef and Schultz, 2009; Ricks et al., 2019). Furthermore, biomolecules other than lipids, such as proteins and glycans, for which metabolic labeling has been well studied (Dieterich et al., 2010; Saxon and Bertozzi, 2000), would in principle also be within the scope of application. We aim to unveil the genetic basis of physiological and pathological lipid metabolism by applying O-ClickFC to primary cells and model animals (Cortez et al., 2020; Fu et al., 2021). Further development of OCDs with better fluorescent properties (e.g., less spectral overlap and fluorogenic abilities) and more specific (sub)organelle membrane localizability would be beneficial to facilitate these future works.

## Materials and Methods

### Key resources table

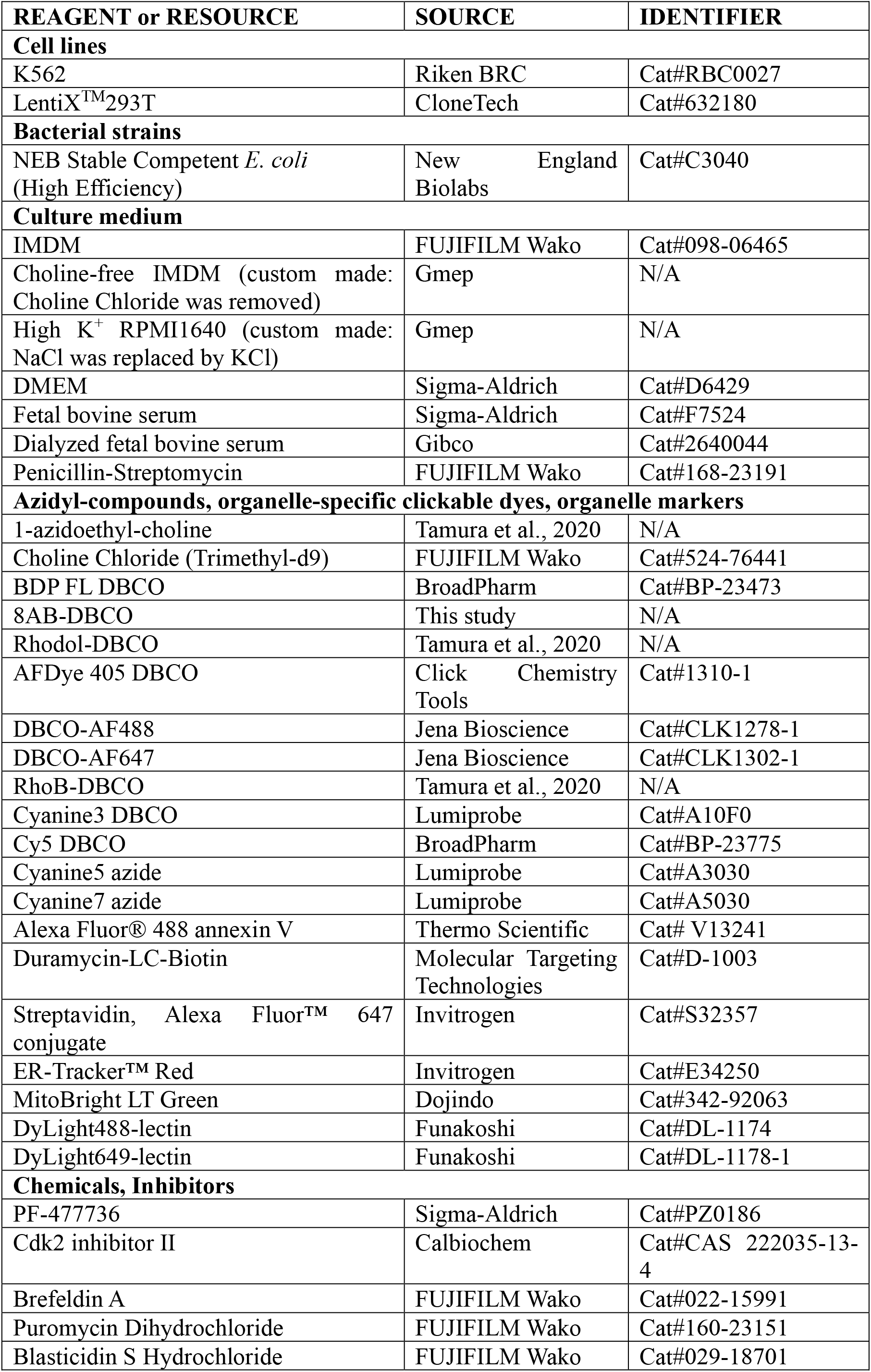
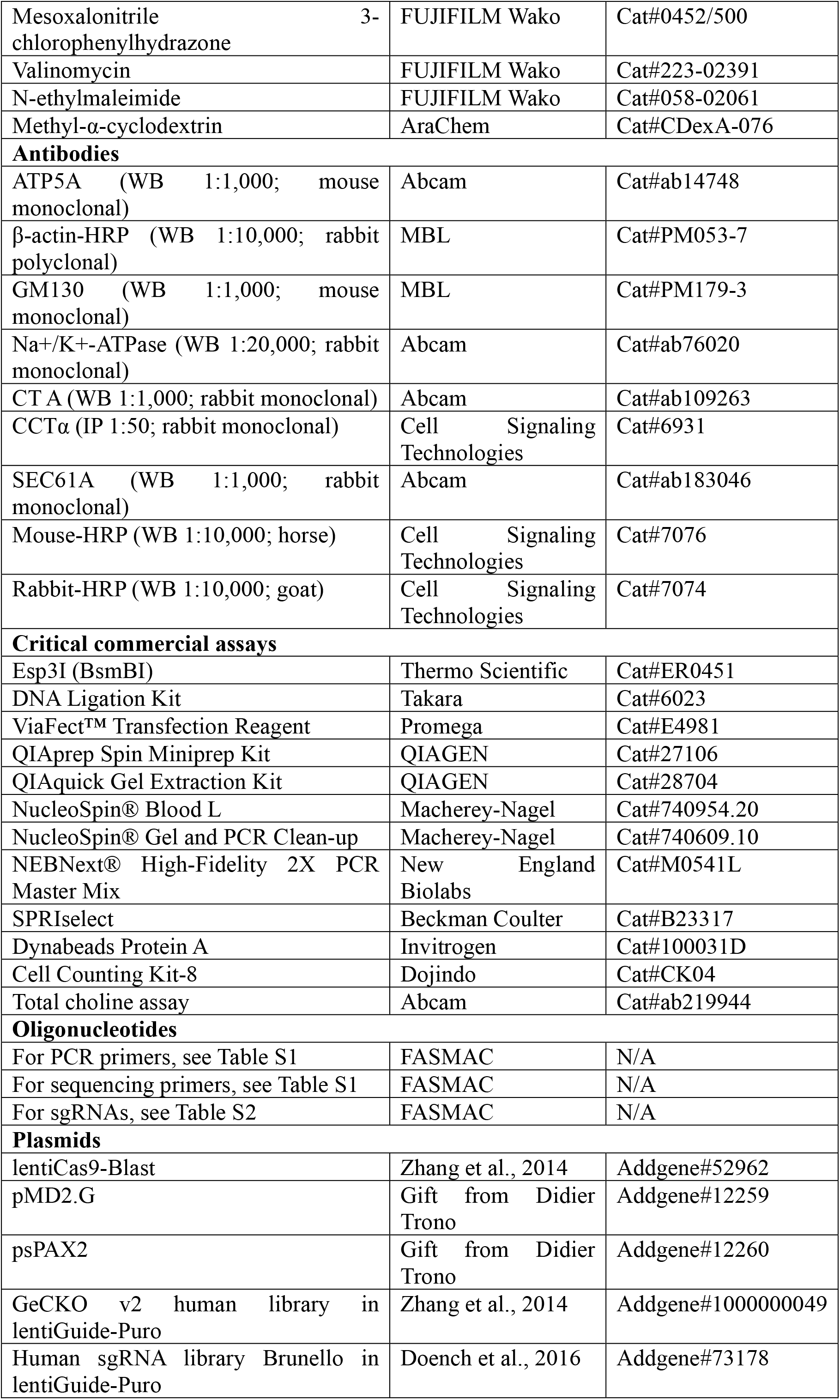
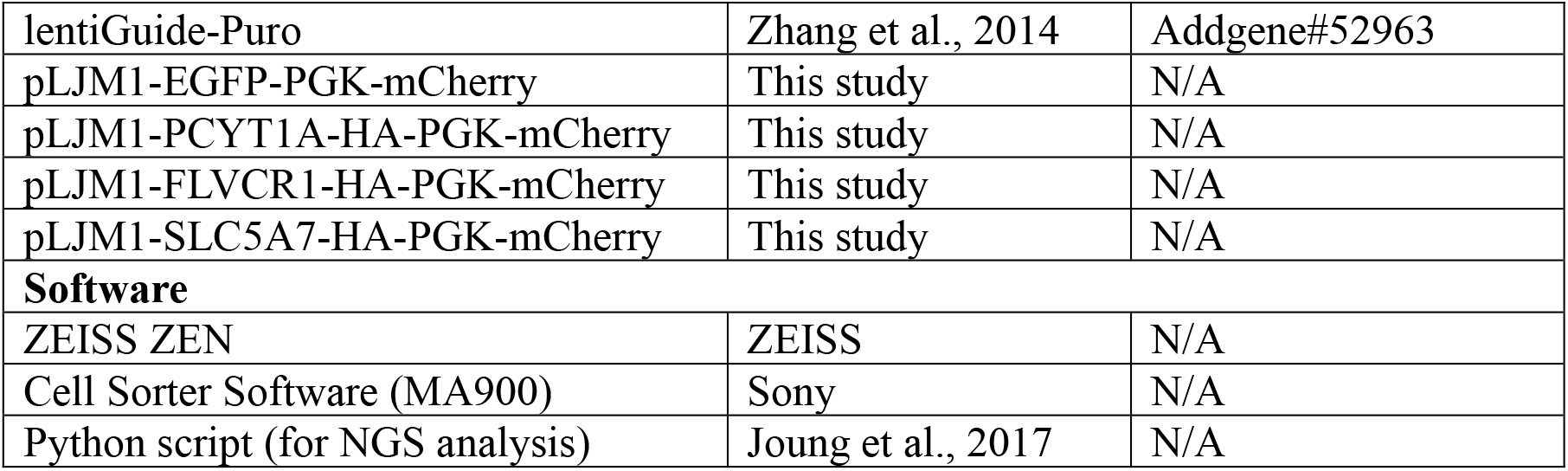

### Experimental model and subject details

#### Cell culture

K562 cells were grown in IMDM containing 10% fetal bovine serum (FBS) and 100 units/mL penicillin-streptomycin (P/S) (hereafter GM). LentiX 293T cells were grown in DMEM containing 10% FBS and 100 U/mL P/S. K562 cells were used in all the assays. LentiX 293T cells were used only for lentivirus production.

### Methods details

#### Synthesis

N_3_-Cho, rhodol-DBCO and rhodamine B-DBCO were prepared as previously described (Tamura et al., 2020). 8AB-DBCO was synthesized according to previous report with minor modifications (Alamudi et al., 2018). Briefly, [2-[(Methylthio)(2H-pyrrol-2-ylidene)methyl]-1H-pyrrole](difluoroborane) (TCI, M2609) was reacted with dibenzocyclooctyne-amine (TCI, A2763). After purification of the crude material, product was confirmed by NMR.

#### Metabolic incorporation of azide-choline

Cells were cultured in Cho-free medium (choline-free IMDM, 10% dialyzed FBS, 100 U/mL P/S) containing 0.1, 1, 10, 100 μM N_3_-Choat 37°C for 5-24 hours depending on the experiments. Most of the data were obtained from the condition of incubation with 10 μM N_3_-Cho for 1 day unless otherwise noted.

#### Organelle-selective labeling of N_3_-PC

After N_3_-Cho incorporation, cell density was adjusted to 1 × 10^6^ cells/mL. Cells were washed with 4% FBS containing IMDM and subsequently incubated with OCDs diluted in 4% FBS/IMDM with the following conditions. ER-Golgi: 100 nM BDP-DBCO, 8AB-DBCO, or Rhodol-DBCO at RT for 15 mins. PM: 100 μM AF405-DBCO, AF488-DBCO, or AF647-DBCO at 15 °C for 30 mins.

Mitochondria: 50 nM RhoB-DBCO, Cy3-DBCO, or Cy5-DBCO at 37 °C for 15 mins, followed by 100 nM Cy5-N_3_ (for quenching the fluorescence signal from unreacted RhoB-DBCO and Cy3-DBCO) or Cy7-N_3_ (for quenching the fluorescence signal from unreacted Cy5-DBCO) at 37 °C for 15 mins, then rinsed three times with a washing medium consisting of high K^+^ RPMI1640, 10% FBS, 50 μM mesoxalonitrile 3-chlorophenylhydrazone, and 20 μM valinomycin. After labeling, cells were washed with GM twice and resuspended in GM to 5-10 × 10^5^ cells/mL for flow cytometry. For ER-Golgi and PM labeling, cells were first labeled with AF405/647-DBCO, and then labeled with 8AB/BDP-DBCO. For ER-Golgi and mitochondrial labeling, cells were first labeled with Cy3-DBCO and then labeled with 8AB-DBCO.

#### Flow cytometric measurements of dye-labeled PC

Flow cytometric analysis of labeled cells was performed using Sony Cell Sorter MA900, equipped with four excitation lasers (488, 405, 561, and 638 nm) and 12-color channels. All four lasers and filters with emission BP 525/50 (Rhodol, BDP, AF488), 585/30 (RhoB, Cy3), 450/50 (8AB, AF405), and 665/30 (Cy5, AF647) were used in this study. For optimal data acquisition, 100 μm sorting chips and following instrument settings were used: FSC threshold value: 5%; Sensor gain: FSC: 3, 525/50: 33.0%, 585/30: 36.0%, 450/50: 39.0%, and 665/30: 40.5%. For each sample, at least 30,000 events were analyzed. The data acquisition, analysis, and image preparation were carried out using the instrument software MA900 Cell Sorter Software (Sony). For FACS-based cell isolation, labeled cells were sorted at 4 °C and collected into GM-containing tubes. After transfer to fresh GM, cells were cultured and subjected to further analysis.

#### Confocal imaging of dye-labeled PC

Labeled cells were deposited on a 35 mm glass-base dish (IWAKI, 11-0609). Imaging of live cells was performed using a Zeiss LSM800 confocal microscope with a Zeiss Plan-Apochromat 100x/1.40 oil objective. The data acquisition, analysis, and image preparation were carried out using the instrument software ZEN (ZEISS). For co-localization analysis, cells labeled with OCDs were stained with the following organelle markers: ER-Tracker Red (ER-Golgi), MitoBright LT Green (mitochondria), or DyLight488/649-lectin (PM).

#### Plasmids

A list of all plasmids and sgRNAs used is provided in Appendix Table S1 and S2 (Excel file). sgRNA-expressing lentiviral plasmids were constructed by inserting 20 bp-long sgRNA oligonucleotide adjacent to the gRNA scaffold in lentiGuide-Puro as previously described (Joung et al., 2017). Briefly, lentiGuide-Puro was digested with BsmBI and 20-bp DNA fragment encoding sgRNA of interest was inserted. Appropriate overhang sequences were appended to each sgRNA sequence for cloning into the lentiGuide-Puro plasmids. For construction of protein-expressing lentiviral plasmids, complementary DNAs (cDNAs) of HA-tagged human PCYT1A, FLVCR1 and SLC5A7 (Uniprot #: P49585-1, Q9Y5Y0-1, and Q9GZV3-1 respectively) were codon optimized and synthesized by AZENTA Inc. Japan. HA-tag epitope was conjugated to the 3’ end of each gene. cDNAs of PCYT1A-HA, FLVCR1-HA and SLC5A7-HA were subcloned into pLJM1-EGFP-PGK-mCherry vector backbone (a derivative of pLJM1-EGFP, Addgene #19319, where *hPGK::mCherry* was inserted) replacing EGFP with the cDNAs.

#### Lentiviral transduction

Lentivirus was prepared according to previous protocol with modification (Joung et al., 2017). LentiX 293T cells were seeded in a 12-well plate and cultured overnight to reach 80-90% confluency. Mixture of 161 μL OptiMEM, 4.84 μL ViaFect, 263.5 ng pMD2.G, 585 ng psPAX2, and 765 ng sgRNA plasmid was added to 1 mL of culture medium. The supernatant was collected 48 hours post-transfection. For lentivirus production of CRISPR-KO library, 1.8 × 10^7^ cells were seeded on a T225 flak in 45 mL of DMEM-based GM and transfected with 15.3 μg pMD2.G, 23.4 μg psPAX2, and 30.6 μg CRISPR library. Lentivirus was condensed to 10x concentration with LentiX Concentrator (CloneTech, 631231).

K562 cells were resuspended in GM containing 10 μg/mL Polybrene and plated at a density of 1.5 × 10^6^ cells/mL x 2 mL/well in 12-well plate. 400 μL of lentiviruses were added to each well, then centrifuged at 1000 x g, 33 °C, 2 hrs (spinfection). For generating stable Cas9-expressing cells, K562 cells were infected with lentiCas9-Blast lentivirus, then cultured and selected with 5 μg/mL blasticidin S. Cas9-expressing K562 cells were infected with lentiGuide-Puro lentivirus. 24 hrs after spinfection, the cells were transferred to a 6-well plate or 10 cm dish, replacing the medium with 0.5 μg/mL Puromycin-containing GM and cultured for 4 days. The medium was replaced every two days. The cells were analyzed five days post-infection.

#### CRISPR screening

CRISPR screening was performed according to previous protocol with modification (Joung et al., 2017). Briefly, lentivirus of GeCKOv2 or Brunello library was used to infect 7.8 × 10^7^ Cas9-expressing K562 cells at an MOI < 0.3, and cells were selected with 0.5 μg/mL puromycin. At four days post-infection, the medium was replaced with the Cho-free medium containing 10 μM N_3_-Cho and incubated at 37 °C for 24 hours. Cells were labeled with BDP-DBCO (ER-Golgi screening), BDP-DBCO & AF405-DBCO (PM screening), or 8AB-DBCO & Cy3-DBCO (mitochondrial screening) as described above. Dead cells were stained with Fixable Viability Dye eFluor 780 (FVD780, ThermoFisher, 65-0865-14) in ER-Golgi and PM screens. Scatter gating (FSC, BSC) was used to remove cell debris, doublet cells and dead cells, and cells with reduced fluorescence intensity in labeled PC were sorted (Figure S4A, S6A, S6E). FACS was performed at a flow rate of 5000-10000 events per second. At least 5 × 10^5^ cells were collected.

#### NGS analysis

Genomic DNA (gDNA) was isolated with NucleoSpin Blood L as the manufacturer’s instructions. For 1^st^ PCR amplification (amplification of lentiviral region in the genome), gDNAs were divided into 16 × 50 μL reactions such that each tube contains 1000-1600 ng of gDNA. Each tube consisted of 5 μL of 5 μM forward and reverse (Lenti-1F and Lenti-2R respectively) primer each, 15 μL of gDNA and water, and 25 μL NEBNext® High-Fidelity 2X PCR Master Mix. PCR cycling conditions: an initial 3 mins at 98 °C; followed by 10 sec at 98 °C, 10 sec at 63 °C, 25 sec at 72 °C for 24 cycles, and a final 2 mins extension at 72 °C. PCR products were purified with NucleoSpin® Gel and PCR Clean-up according to the manufacturer’s instructions. For 2^nd^ PCR amplification (amplification of sgRNAs), 150-170 μL of purified PCR products were divided into 24 × 50 μL reactions. Each tube consisted of 5 μL of 5 μM forward primers (Mixture of NGS-Lib-Fwd-1 to -10), 5 μL of 5 μM uniquely-barcoded reverse primer (from NGS-Lib-KO-Rev-MT1 to 8), 15 μL of PCR product and water, and 25 μL NEBNext® High-Fidelity 2X PCR Master Mix. PCR cycling conditions: an initial 3 mins at 98 °C; followed by 10 sec at 98° C, 10 sec at 63 °C, 25 sec at 72 °C for 8 cycles, and a final 2 mins extension at 72 °C. PCR products were purified with NucleoSpin Gel and PCR Clean-up. Amplicons (250-270 bp) were verified by 2% agarose gel electrophoresis. Samples were further purified by x0.6/1.2 double size selection using SPRIselect (Beckman Coulter, B23317), then sequenced on a Nextseq500 (Illumina), loaded with a 20% spike-in of PhiX DNA. Multiple screens were performed (ER-Golgi: two times (GeCKOv2 and Brunello); PM: one time (Brunello); mitochondria: two times (Brunello)). NGS results were analyzed according to previous Python script. The sgRNA fold change was determined for generating the gene rank data. Candidate genes in the top 100 or 200 were picked up for further verification Selected genes and sgRNAs were summarized in Table S2.

#### Individual testing of candidate genes

Cas9-expressing K562 cells were infected with 400 μL of sgRNA lentivirus (lentiGuide-Puro, Table S2) per well as described above. Cells were then cultured with 0.5 μg/mL puromycin for four days. sgControl-transduced cells were stained with 1 μM CPM (7-Diethylamino-3-(4’-Maleimidylphenyl)-4-Methylcoumarin, ThermoFisher, used for cell identification in flow cytometric assay) diluted in serum-free IMDM at 37 °C for 15 mins, and washed with GM twice. CPM-labeled (nontargeting control) and unlabeled (CRISPR-KO) cells were then mixed in 1:1 ratio, and cultured in 10 μM N_3_- Cho containing Cho-free medium at 1 × 10^6^ cells/mL density for 24 hrs. On the following day, cells were labeled with OCDs, then analyzed with the flow cytometer. For confirmation of OPM labeling in TMEM30A-KO cells, AF405-DBCO was also used without CPM labeling. Median fluorescence intensity (MFI) was used to compare N_3_-PC levels between control and CRISPR-KO cells. Two or more distinct sgRNAs targeting the same gene that displayed <80% ER-Golgi N_3_-PC for ER-Golgi screen, >60% ER-Golgi N_3_-PC and <80% PM N_3_-PC for PM screen, or >60% ER-Golgi and <80% mitochondrial N_3_-PC for the mitochondrial screen compared to nontargeting control sgRNA were considered as hits. Protein functions of hit genes were described according to UniProt database.

#### FLVCR1-KO rescue assay

Cas9-expressing K562 cells were infected with sgControl or sgFLVCR1 lentivirus as described above. After four days post-transduction, FLVCR1-KO cells were incubated with N_3_-Cho for 1 day, labeled with BDP-DBCO, and cells with fluorescence intensities between 10^3^ and 10^4^ (dim population) were isolated by FACS. Control cells were infected with lentivirus expressing mCherry and GFP, and the isolated FLVCR1-KO cells were infected with lentivirus expressing mCherry together with GFP, PCYT1-HA, FLVCR1-HA, or SLC5A7-HA. After three days culture, cells were again treated with N_3_-Cho for 1 day and labeled with 8AB-DBCO. mCherry-positive cell population (MFI of mCherry >10^3^) was subjected to flow cytometric measurement of 8AB-labeled PC. For bulk-cell analysis of FLVCR1-KO subpopulation with recovery in N_3_-PC labeling, mCherry-positive and 8AB-positive cells (MFI: mCherry >10^3^, 8AB >10^4^) were isolated by FACS and subjected to total choline assay and cell growth assay.

#### LC-MS/MS analysis of phospholipids

For PC-D9 analysis, cells were incubated in Cho-free medium containing 10 μM ChoD9 at 37 °C for 24 hrs at 1 × 10^6^ cells/mL. Lipids were extracted by the Bligh & Dyer method and were subjected to LC-MS/MS analysis using LC-30AD HPLC system (Shimadzu) coupled to triple-quadrupole LCMS-8040 mass spectrometer (Shimadzu) as previously described (Tamura et al., 2020). The precursor ions of m/z 184 and 193 were monitored for detection of PC and PC-D9 (positive ion mode), respectively. The total ion current chromatogram of PC and PC-D9 was recorded to quantify PC-D9 content (presented as percentage of PC-D9 in total of PC and PC-D9). For extraction of phospholipids in the outer leaflet of the plasma membrane (Li et al., 2016), 1 × 10^7^ cells were incubated in PBS containing 40 mM methyl-α-cyclodextrin (AraChem) and 1.5 mM 1,2-dioleoyl-sn-glycero-3-phospho-(1’-rac-glycerol) (DOPG, Avanti) for 1 hour at RT. After centrifuging, the supernatant was collected and subjected to lipid extraction and LC-MS/MS analysis. PE was detected by neutral loss scanning of m/z 141 (positive ion mode). Amount of PC or PE was quantified as total intensities of choline-or ethanolamine-containing lipid species detected in the range of m/z 590-940, respectively.

#### LC-MS/MS analysis of phosphorylated PCYT1A

1 × 10^6^ K562 cells cultured in GM were collected and lysed with Cell Extraction Buffer (Invitrogen, FNN0011) with protease inhibitor (1x protease inhibitor complete, 1mM PMSF). PCYT1A was immunoprecipitated according to the manufacturer’s instruction using 100 μL Dynabeads Protein A with 2 μg of anti-CCTA antibody (CST) per purification. Beads were resuspended in 100 μL of 100 mM Tris pH 8.0 buffer. 500 ng of Trypsin/Lys-C Mix was added and incubated at 37 °C O/N, centrifuged and the supernatant was transferred to new tubes. Peptides were reduced by DTT, alkylated by iodoacetic acid. Alkylation was stopped by cysteine, then desalted by C18 ZipTip. Peptides were then subjected to nanoLC-MS analysis using the UltiMate 3000 RSLCnano LC System (ThermoFisher). Mobile phase A: 0.1% formic acid and B: 0.1% formic acid in 80% acetonitrile. Mobile phase gradient: 2% B at 0 min, 5% B at 4 min, 32% B at 74 min, and 65% B at 84 min. Flow rate: 750 nl/min at 0–4 min, 200 nl/min at 4–84 min. Peptide eluents were analyzed by Q Exactive HF-X (Thermo Scientific). Acquired data were processed with PEAKS Studio (Bioinformatics Solutions Inc). The phosphorylation sites identified in this analysis were compared with the UniProt database (ID: P49585).

#### Pharmacological treatment

For inhibition of CHK1 and CDK2, K562 cells were cultured in Cho-free medium at 1.0 × 10^6^ cells/mL density with 10 μM N_3_-Cho and following inhibitors at the given concentration at 37 °C for 24 hrs. CHK1 inhibitor: 1 μM PF-477736; CDK2 inhibitor: 10 μM Cdk2 inhibitors II. Cells were then labeled with BDP-DBCO and subjected to flow cytometric analysis. For PC-D9 quantification, cells were first cultured with inhibitors in Cho-free medium at 37 °C for 24 hours, and then treated with ChoD9, followed by LC-MS/MS analysis. For inhibition of PM trafficking, cells were first cultured in Cho-free medium at 1.0 × 10^6^ cells/mL O/N, then treated with 10 μM N_3_-Cho and 10 μM Brefeldin A (BFA) at 37 °C for 5 hrs. After the incubation, cells were labeled with OCDs, then analyzed with flow cytometry and microscopy.

#### Flow cytometric analysis of cell surface PS and PE

For assessing PS on cell surface, cells were washed with Annexin binding buffer (Invitrogen, V13246) once, labeled with AF488-Annexin V diluted in Annexin buffer (1:20) at RT for 15 mins, followed by a one-time wash with Annexin buffer, then analyzed with flow cytometry. For assessing PE, cells were washed with 0.5% fatty acid (FA)-free BSA/PBS (Wako) once, labeled with 45 μM Duramycin-LC-Biotin/Strepavidin-647 pre-conjugated complex (1:50) diluted in 0.5% Fatty Acid-free BSA/PBS at 15°C for 15 mins, followed by a one-time wash with 0.5% FA-free/PBS, then analyzed with flow cytometry.

#### Cell growth assay

2500 cells were inoculated on a 96-well plate per well and cultured for 24-48 hours. Cell Counting Kit-8 was added into each well (10 ul per 100 ul medium) and incubated at 37°C for 1 hour. The absorbance at 450 nm was measured with a plate reader (TECAN, Infinite 200 PRO).

#### GPMV isolation

GPMVs were prepared according to previous study with modification (Sezgin et al., 2012). Briefly, ChoD9 treated K562 cells were washed with GPMV buffer (10 mM HEPES, 150 mM NaCl, 2 mM CaCl_2_, pH 7.4), then incubated in 2 mM N-ethylmaleimide/GPMV buffer at 37 °C for 1 hour allowing the cells to produce a sufficient amount of GPMVs. Supernatants containing GPMVs were collected, centrifuged at 1,000 x g for 2 mins to remove cells, followed by 16,000 x g for 10 mins to sediment GPMVs. Collected GPMVs were subjected to western blotting or LC-MS/MS analyses.

#### SDS-PAGE and western blotting

K562 cells or GPMVs were lysed in Cell Extraction Buffer with protease inhibitor (1x protease inhibitor complete, 1 mM PMSF). 20 μg of lysate was run on SDS-PAGE gels (BIO-RAD, 4561095) or on Phos-Tag SDS-PAGE gel (10%, Wako, 193-16711). Phos-Tag gels were incubated in transfer buffer containing 1 mM EDTA for 10 min, and then soaked in normal transfer buffer for 10 min. Proteins were transferred to a PVDF membrane and standard western blotting procedures were subsequently followed.

#### Total choline assay

Total choline assay was performed according to the manufacturer’s instruction. Briefly, 5 × 10^5^ cells were lysed with 200 μL of the provided Assay Buffer, and 2 × 50 μL were used for the assay in a 96-well plate. 50 μL Choline reaction mix was added into each sample, then incubated at RT for 30 mins in the dark. The amount of Choline was quantitated with a fluorescence microplate reader (TECAN, Infinite® 200 PRO) at Ex./Em. = 540/590 nm.

#### Statistical analysis

Bar graphs were represented as mean ± SEM. Heatmaps and dot plots in individual testing of genes and sgRNAs were generated by taking mean of the data. Number of data point per group was indicated in the figures or figure legends. Statistical analyses were performed using a two-way unpaired t-test.

## Acknowledgments

The authors would like thank K. Uchida, A. Fujisawa, K. Kunieda, and H. Nonaka for chemical synthesis; and T. Kondo, Y Sando, and the NGS Core Facility of the Kyoto University Graduate Schools of Biostudies for NGS analysis. MS analysis of protein was performed at the Kazusa DNA Research Institute. We thank Edanz (https://jp.edanz.com/ac) for editing a draft of this manuscript. This work was supported by a Grant-in-Aid for Scientific Research on Innovative Areas “Chemistry for Multimolecular Crowding Biosystems” (JSPS KAKENHI Grant 17H06348), the Japan Science and Technology Agency (JST) ERATO Grant JPMJER1802 to I.H.; a Grant-in-Aid for Scientific Research on Innovative Areas ‘Integrated Bio-metal Science’ (19H05764), Scientific Research (B) (21H02058) to T.T.; and a Grant-in-Aid for Early-Career Scientists (19K16075), JST PRESTO Grant JPMJPR20EA to M.T.

## Author contributions

M.T., T.T., and I.H. conceived the project and designed the experiments. M.T. and N.T. performed the experiments and data analysis. K.N. supervised the MS analysis of lipid. M.T., N.T., T.T., and I.H. wrote the manuscript with input from all authors.

---

**Figure S1.**
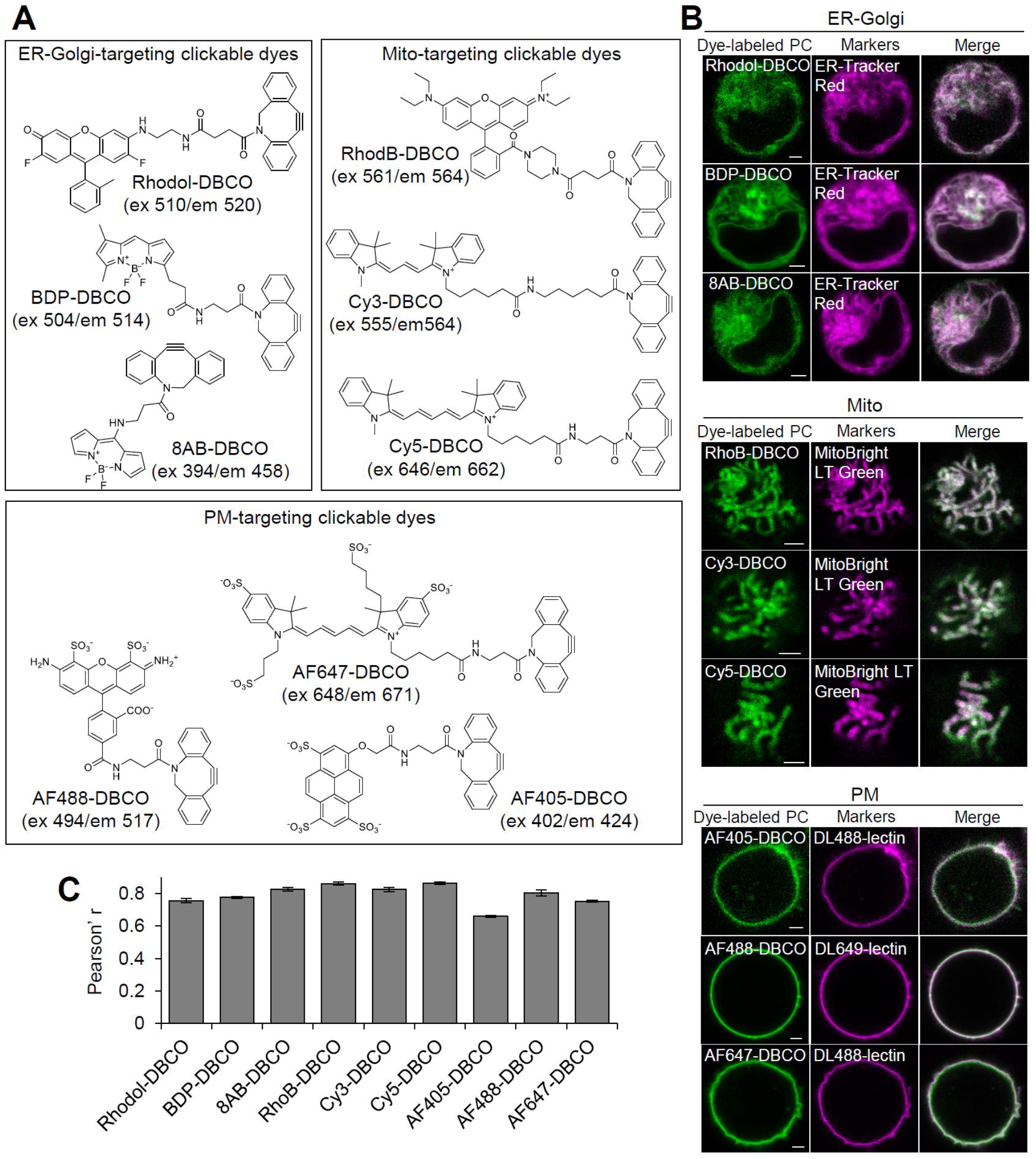
Supporting data for imaging of organelle-selective N_3_-PC labeling, related to Figure 2. (A) Chemical structures of organelle-targeting clickable dyes (OCDs). Rhodol, RhodB, and 8AB were prepared in-house, and the others were obtained commercially. (B) Confocal images of K562 cells treated with N_3_-Cho and labeled with OCDs (ER-Golgi: Rhodol-DBCO, BDP-DBCO, 8AB-DBCO; PM: AF405-DBCO, AF488-DBCO, AF647-DBCO; Mitochondria: RhoB-DBCO, Cy3-DBCO, Cy5-DBCO) and organelle markers (ER-Golgi: ER-Tracker Red; PM: DL488/649-lectin; Mitochondria: MitoBrightLT Green). Scale bar = 2 μm. (C) Graph shows the Pearson’s correlation coefficient from each group (n = 4–8). Bar graphs represent mean ± SEM.

**Figure S2.**
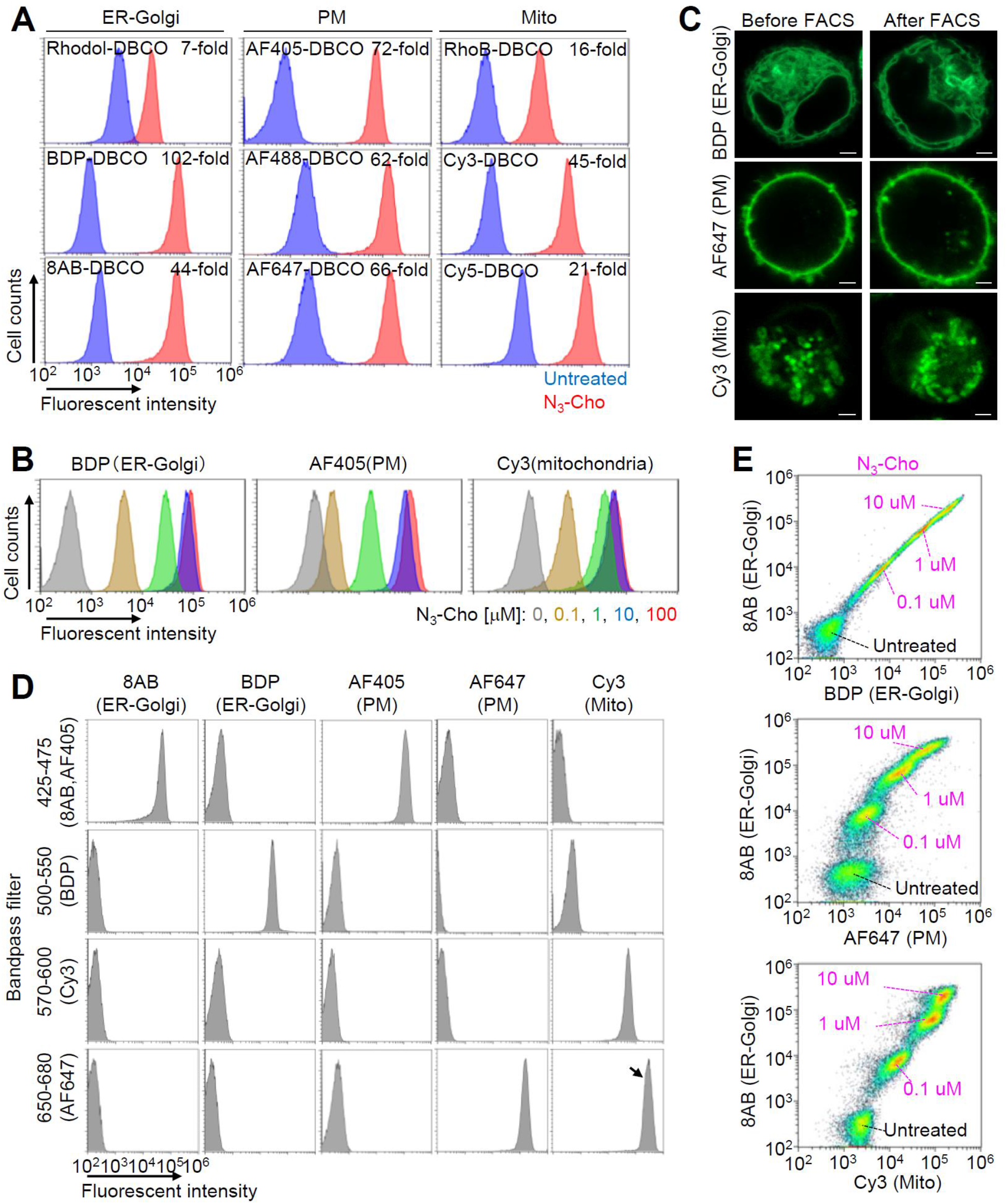
Supporting data for flow cytometric measurements of organelle-selective N_3_-PC labeling, related to Figure 2. (A) Quantification of organelle-selective N_3_-PC labeling. K562 cells incubated overnight with 0 (control) or 10 μM N_3_-Cho were labeled with OCDs and analyzed by flow cytometry. MFIs were used to calculate fold change between control and N_3_-Cho-treated cells (n = 2–3). (B) Flow cytometric analyses of K562 cells treated with varying concentrations of N_3_-Cho (0, 0.1, 1, 10, 100 μM) overnight, and labeled with BDP-DBCO, AF405-DBCO, or Cy3-DBCO. Overlaid histograms show that fluorescence increase of labeled PC depended on the N_3_- Cho concentration and saturated at 10 uM. (C) Images of PC labeled with BDP-DBCO, AF647-DBCO, or Cy3-DBCO before (left) and after (right) FACS, showing no major changes in fluorescence intensity or intracellular distribution. Scale bar = 2 μm. (D) Test of fluorescence leakage in flow cytometric analysis of OCDs with multiple different bandpass filters. Histograms show no obvious crossover in the following combinations: 8AB-BDP, 8AB-Cy3, 8AB-AF647, BDP-AF405, BDP-AF647, and AF405-Cy3. The arrow denotes fluorescence from quencher Cy5-N_3_. (E) Two-dimensional flow cytometric analysis of N_3_-PC distribution with dual organelle labeling. K562 cells were separately treated with different concentrations of N_3_-Cho (0, 0.1, 1, 10 μM) overnight, and pooled into one tube. After dual labeling with 8AB-DBCO and BDP-DBCO, 8AB-DBCO and AF647-DBCO, or 8AB-DBCO and Cy3-DBCO, cells were analyzed by flow cytometry. Dot plots show that a bulk population was separated into four groups with different fluorescence levels corresponding to N_3_-Cho concentrations.

**Figure S3.**
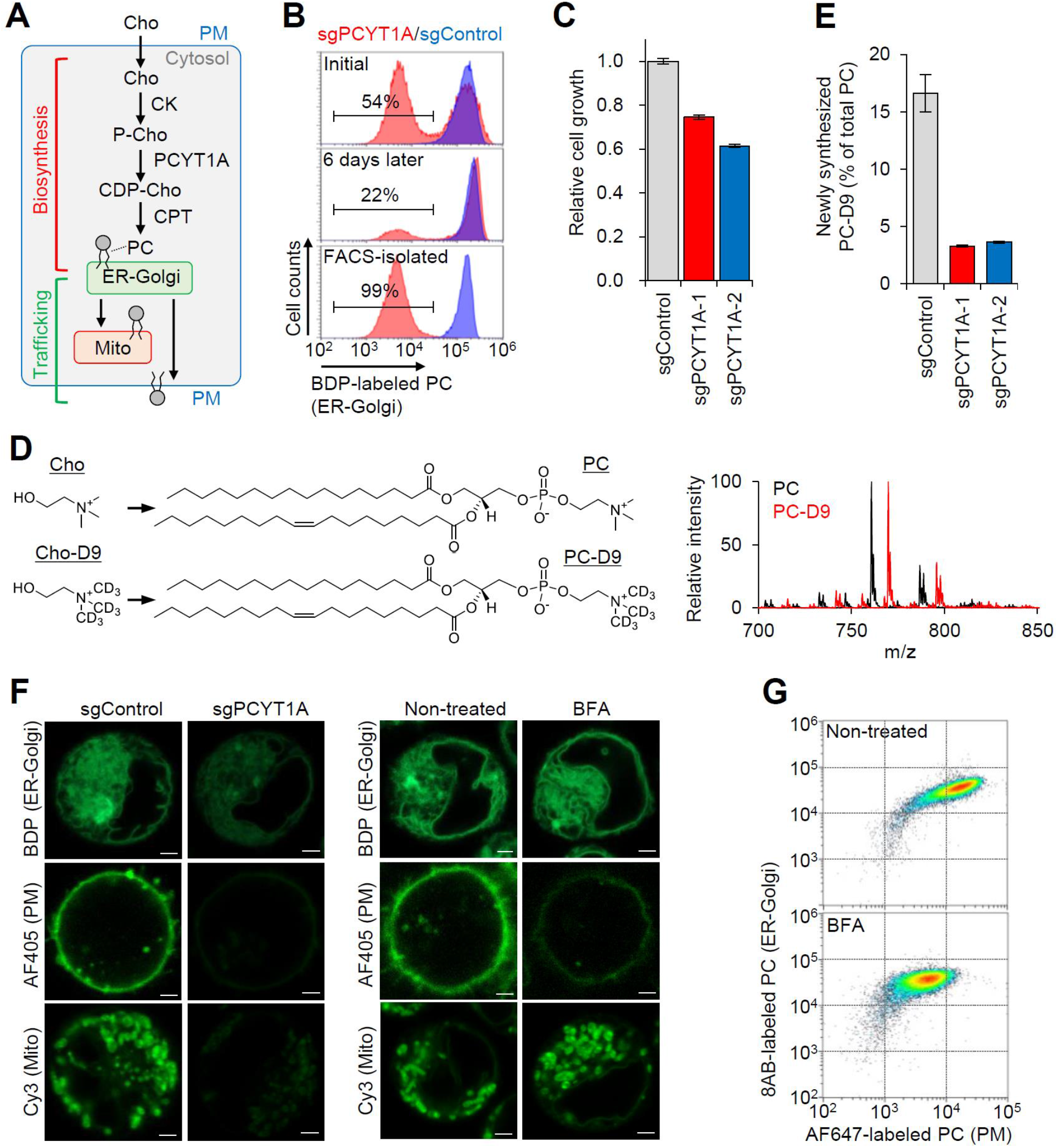
Supporting data for characterization of PCYT1A-deficient cells and BFA-treated cells, related to Figure 2. (A) Schematic of PC biosynthesis and trafficking in mammalian cells. (B) FACS isolation of PCYT1A-deficient cells based on N_3_-PC level in ER-Golgi. In sgPCYT1A-transduced cells, the BDP-dim population (fluorescence intensity between 103 and 104) decreased over time (initial vs 6 days later) and was isolated by FACS. Sorted cells were expanded and subjected to a second analysis (FACS-isolated). (C) Proliferation of *PCYT1A*-KO K562 cell (n = 4). (D) LC-MS/MS analysis of *de novo*-synthesized PC-D9 in whole-cell lysates of *PCYT1A*-KO K562 cells. A precursor ion scan of m/z 184 and 193 was used for PC and PC-D9, respectively. (E) Quantification in (D) (n = 3). PC-D9 content was calculated as the percentage of PC-D9 to total PC and PC-D9. (F) Confocal images of control (left), *PCYT1A*-KO cells (middle-left), non-treated (middle-right), and BFA-treated cells (right) labeled with different OCDs. Cells prepared for flow cytometry in Figure 2D and E were subjected to confocal laser scanning microscopy. All three types of fluorescence (BDP, AF405, and Cy3) were decreased in *PCYT1A*-KO cells, compared with control CRISPR cells. No major changes in ER-Golgi or mitochondrial fluorescence were observed, but a decrease in PM fluorescence was observed in BFA-treated cells compared with non-treated cells. Scale bar = 2 μm. (G) Two-dimensional flow cytometric analysis of N_3_-PC distribution in BFA-treated cells. K562 cells were treated with N_3_-Cho in the absence (top) or presence (bottom) of BFA, and then labeled with 8AB-DBCO and AF647-DBCO. MFI of AF647 was decreased, but 8AB was unchanged in BFA-treated cells compared with non-treated cells. Bar graphs represent mean ± SEM.

**Figure S4.**
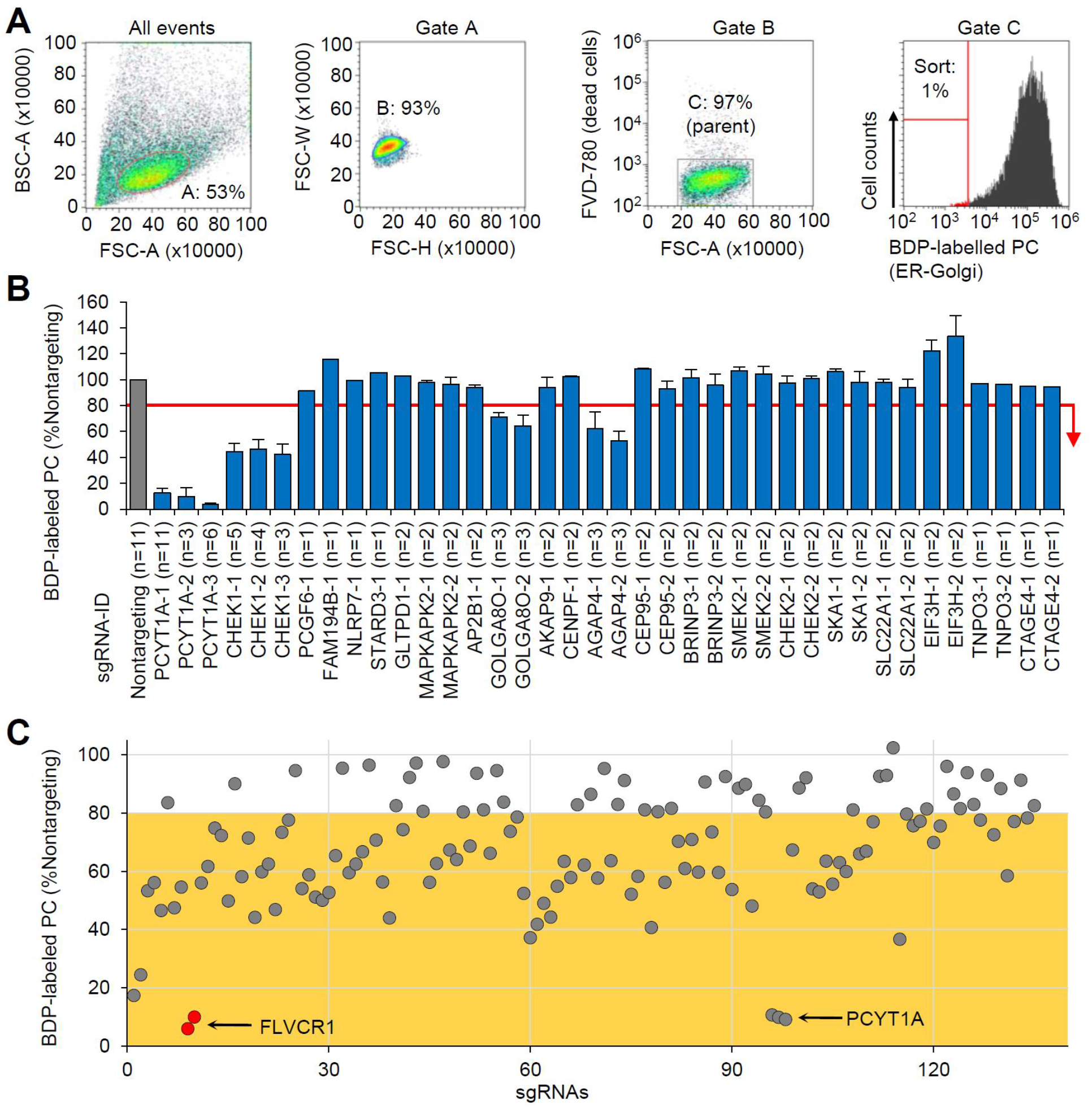
Supporting data for CRISPR-KO screening of N_3_-PC in the ER-Golgi, related to Figure 3. (A) FACS gating strategy for ER-Golgi-focused screening. Using forward scatter (FSC) and back scatter (BSC), live cells were distinguished from cell debris and dead cells (All events). Singlet cells were separated from doublet cells on physical parameters (FSC-H vs FSC-W, Gate A). Live cells were confirmed by negative staining with the dead-cell marker FVD780 (Gate B). Cells exhibiting weak fluorescence indicating BDP-labeled PC (∼1% of parent population) were sorted (Gate C). (B) Individual testing of 38 sgRNAs targeting 22 genes selected from the top 100 in the GeCKOv2 screening. MFI was used to compare ER-Golgi N_3_-PC levels between nontargeting and gene-targeting sgRNAs. Cells that exhibited lower than 80% fluorescence intensity compared with nontargeting control were considered hits. The 80% threshold is shown as a red line in the graph. Bar graphs represent mean ± SEM. (C) Individual testing of 135 sgRNAs targeting 77 genes selected from the top 100 in the Brunello screening. Dot plot shows 90 sgRNAs below the threshold level of 80% (orange area). sgRNAs targeting *FLVCR1* and *PCYT1A* are highlighted. Dot plot represents the mean.

**Figure S5.**
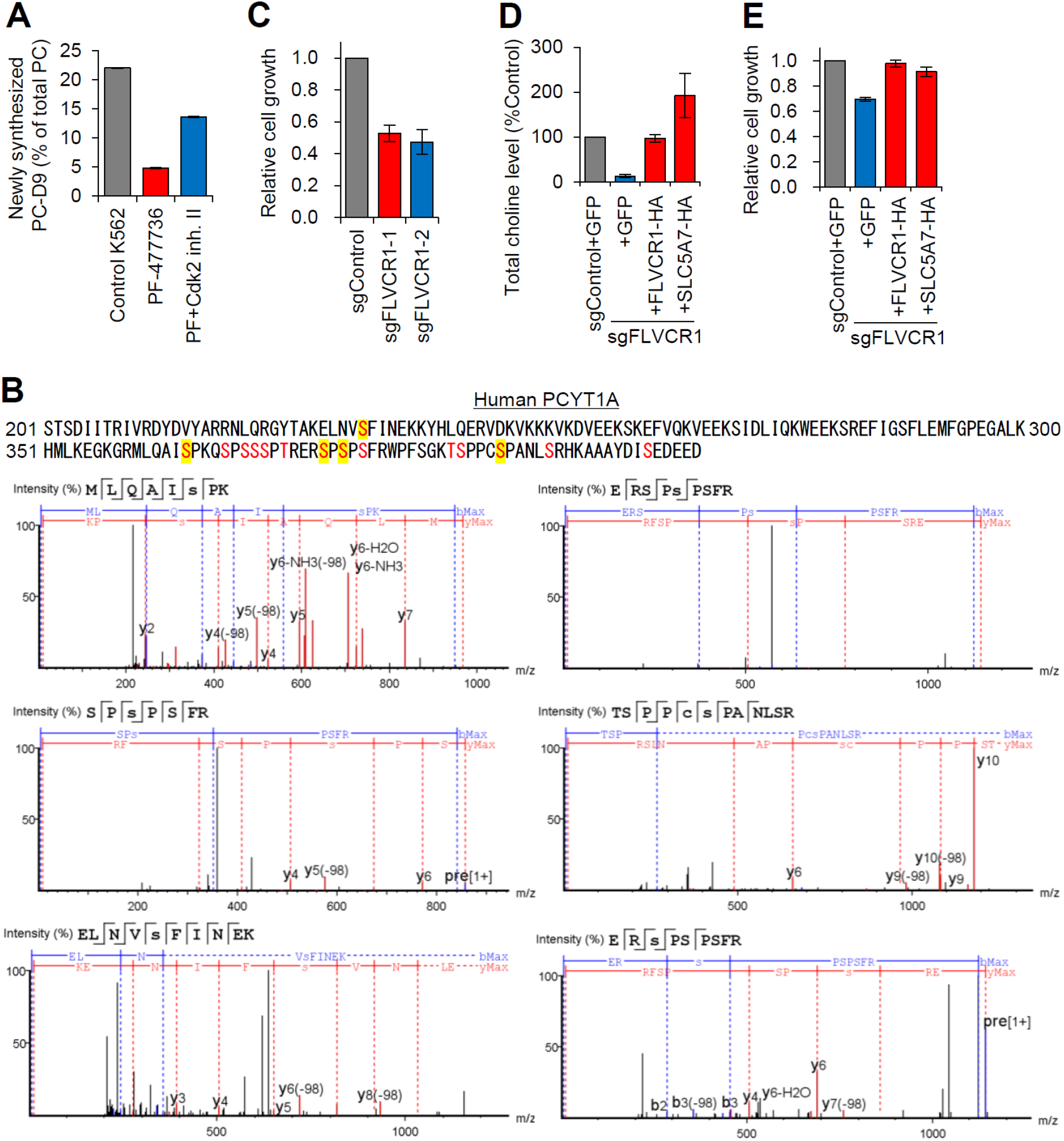
Supporting data for roles of *CHEK1* and *FLVCR1* in PC synthesis, related to Figure 4. (A) Quantitative LC-MS/MS analysis of PC-D9 levels in whole-cell lysate of K562 cells treated with CHK1i (PF-477736) and CDK2i (Cdk2 inhibitor II) (n = 3). (B) Phosphorylation sites of PCYT1A. Phosphorylation of serine and threonine in red characters was previously reported based on sequence similarity (UniProtKB reference number: P49585). Phosphorylated serine residues highlighted in yellow were detected in this MS/MS analysis of PCYT1A protein, which was extracted from K562 cells cultured in normal growth medium. (C) Proliferation of *FLVCR1*-KO K562 cells (n = 3). (D) Total endogenous choline levels in control or *FLVCR1*-KO K562 cells expressing GFP, HA-tagged FLVCR1, or HA-tagged SLC5A7 (n = 4). (E) Proliferation of control or *FLVCR1*-KO K562 cells expressing GFP, HA-tagged FLVCR1, or HA-tagged SLC5A7 (n = 3). Bar graphs represent mean ± SEM.

**Figure S6.**
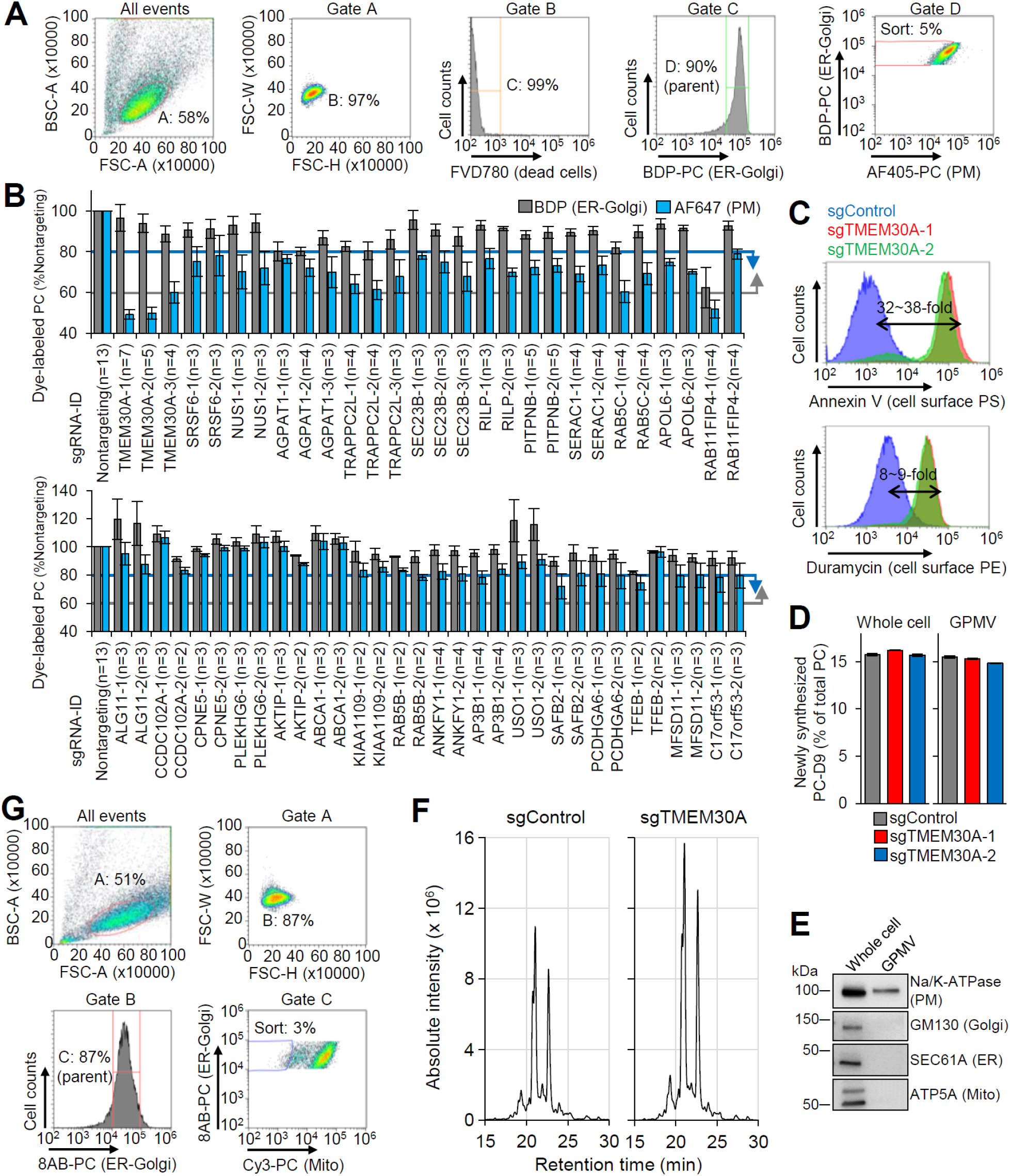
Supporting data for CRISPR-KO screening focusing on subcellular PC trafficking of N_3_-PC, related to Figure 5. (A) FACS gating strategy in PM-focused screening. Live singlet cells were obtained at Gate B. The majority of cells with robust BDP fluorescence intensity indicating ER-Golgi N_3_-PC were separated (Gate C). Cells exhibiting weak AF405 fluorescence in PM N_3_-PC (∼5% of the parent population) were sorted (Gate D). (B) Individual testing of 60 sgRNAs targeting 28 genes selected from the top 100. MFIs of labeled PC were compared between nontargeting and gene-targeting sgRNAs. The upper graph shows 28 sgRNAs satisfying the criteria: > 60% ER-Golgi labeling (grey line) and < 80% PM labeling (blue line). The lower graph shows the remaining 32 sgRNAs that failed to fulfill the criteria. (C) FACS analysis of cell-surface PS and PE with PS-binding annexin V and PE-binding duramycin (n = 3). (D) LC-MS/MS analysis of PC-D9 contents in whole-cell lysate and giant plasma membrane vesicle (GPMV) from control and *TMEM30A*-KO cells (n = 3). (E) The purity of GPMV was confirmed by WB. (F) LC-MS/MS analysis of PE extracted from the PM outer leaflet of control and *TMEM30A*-KO cells by methyl-α-cyclodextrin-mediated lipid exchange. PE was increased in the outer PM of *TMEM30A*-KO cells. (G) FACS gating strategy in mitochondrial screening. Live singlet cells were obtained at Gate A. The majority of cells with robust 8AB fluorescence intensity in ER-Golgi N_3_-PC were separated (Gate B). Cells exhibiting weak Cy3 fluorescence in mitochondrial N_3_-PC (∼3% of the parent population) were sorted (Gate C). All bar graphs represent mean ± SEM.

